# 5-hydroxymethylcytosine deposition mediates Polycomb Repressive Complex 2 function in *MYCN*-amplified neuroblastoma

**DOI:** 10.1101/2025.07.05.663256

**Authors:** Mohansrinivas Chennakesavalu, Gabriel L. Lopez, Varsha Gupta, Kelley Moore, Yuqing Xue, Josie Majowka, Sahil Veeravalli, Ryan Borchert, Monica Pomaville, Alexandre Chlenski, Chuan He, Andrea Piunti, Mark A. Applebaum

## Abstract

*MYCN*-amplification is a strong predictor of poor prognosis in neuroblastoma, an embryonal malignancy that accounts for 15% of pediatric cancer deaths. Here, we found that *MYCN*-amplified neuroblastoma tumors had increased 5-hydroxymethylcytosine (5-hmC) deposition on Polycomb Repressive Complex 2 (PRC2) target genes. 5-hmC and H3K27me3, a catalytic product of PRC2, directly co-localized at the nucleosomal level in *MYCN*-amplified neuroblastoma. Genes with co-localization of 5-hmC/H3K27me3 were involved in development related pathways and were transcriptionally repressed in *MYCN*-amplified neuroblastoma. Inhibition of 5-hmC deposition resulted in a loss of H3K27me3 on protein-coding genes and sensitized neuroblastoma to DNA demethylating agents. 5-hmC deposition predisposed H3K27me3 marked genes to transcriptional activation upon PRC2 inhibition with tazemetostat. Low expression of genes marked by 5-hmC/H3K27me3 was associated with poor clinical outcome. Our results suggest that 5-hmC/H3K27me3 co-operate to repress mediators of development highlighting a novel link between DNA and chromatin modifications with potential therapeutic implications in *MYCN*-amplified neuroblastoma.

## INTRODUCTION

Neuroblastoma is an embryonal malignancy that accounts for nearly 15% of pediatric cancer deaths^1,2^. Neuroblastoma originates from cells of the neural crest, and the onset of this disease is thought to represent abnormal development of the peripheral nervous system^3^. Neuroblastoma is a biologically heterogeneous disease with a range of clinical manifestations. Patients with low-risk (LR) and intermediate-risk (IR) disease have 5-year survival rates of over 90% and LR patients often experience spontaneous regression, while patients with high-risk (HR) disease have a 5-year survival rate of approximately 60% despite intensive multimodal therapeutic approaches^4^. *MYCN-* amplification is a strong predictor of poor prognosis and is an established driver of approximately half of HR neuroblastoma tumors, while *MYC* overexpression is a driver in an additional 20% of HR tumors^5^.

Unlike most cancers of adults, somatic mutations are infrequent in neuroblastoma and epigenetic aberrations are key drivers of oncogenesis^6–8^. Through CRISPR-Cas9 screening, *MYCN-*amplified neuroblastoma was found to be selectively dependent on Polycomb Repressive Complex 2 (PRC2)^9,10^. PRC2 is a multiprotein complex formed by a core composed of EZH2 (or its homolog EZH1), SUZ12, EED and RBBP4/7. Through the methyltransferase activity of the EZH2 subunit, PRC2 catalyzes the mono-, di-, and tri-methylation of lysine 27 of histone 3 (H3K27me1-2-3). H3K27me3 is a histone mark associated with closed chromatin and transcriptional repression^11,12^. PRC2 is involved in a variety of different biological processes and, most importantly, its deregulation is the crucial driving event in several pediatric cancers^12,13^. In neuroblastoma, expression of MYCN and MYC target genes were found to be inversely correlated with the expression of PRC2 target genes. Further, pharmacologic and genetic inactivation of PRC2 induced neuronal differentiation and suppressed in-vitro and in-vivo neuroblastoma tumor growth^9,10^.

In contrast to H3K27me3, 5-hydroxymethylcytosine (5-hmC) generally marks open chromatin and is associated with active gene expression in several cancers, including neuroblastoma^14,15^. 5-hmC is a stable intermediate generated in the process of the demethylation of 5-methylcytosine (5-mC), a process catalyzed by the TET family of proteins^16^. We previously showed the prognostic utility of genome-wide 5-hmC profiling in both tumors and circulating cell free DNA (cfDNA) isolated from patients with neuroblastoma^15,17^. Unexpectedly, we found that PRC2 target genes, which should be transcriptionally repressed, had higher 5-hmC deposition in HR compared to LR tumors and in cfDNA from patients with metastatic and relapsed disease^15,17^. Interestingly, previous studies have shown that 5-hmC is disproportionately enriched at promoters of PRC2 repressed regulators of development in mouse embryonic stem cells (mESCs), but not in more differentiated cells^18–21^. Thus, we hypothesized a cooperative function between 5-hmC and PRC2 at transcriptionally repressed genes that mediates an undifferentiated state in *MYCN*-amplified neuroblastoma.

## RESULTS

### MYCN-amplified tumors have increased 5-hmC on PRC2 target genes

To examine the relationship between 5-hmC deposition and *MYCN*-amplification in neuroblastoma, we compared genome-wide 5-hmC profiles from neuroblastoma derived diagnostic tumor biopsy samples with (n=22) and without (n=83) *MYCN*-amplification (dbGaP accession: phs001831.v1.p1). Principal Component Analysis (PCA) revealed that principal component 2 correlated to *MYCN-*amplification status and explained 14.13% of the variation among tumor samples (**FIGURE 1A**). We identified 4,509 genes with significantly different 5-hmC deposition (FDR<0.05) between *MYCN-* amplified (n=22) and non-amplified (n=83) tumors (**FIGURE 1B**). Unsupervised clustering of 105 diagnostic tumor samples using the 901 genes with the most differential 5-hmC deposition from this analysis clearly distinguished tumor samples with and without *MYCN*-amplification (**SUPPLEMENTAL FIGURE 1**), highlighting the specificity of this gene set to *MYCN*-amplification status.

**FIGURE 1:**
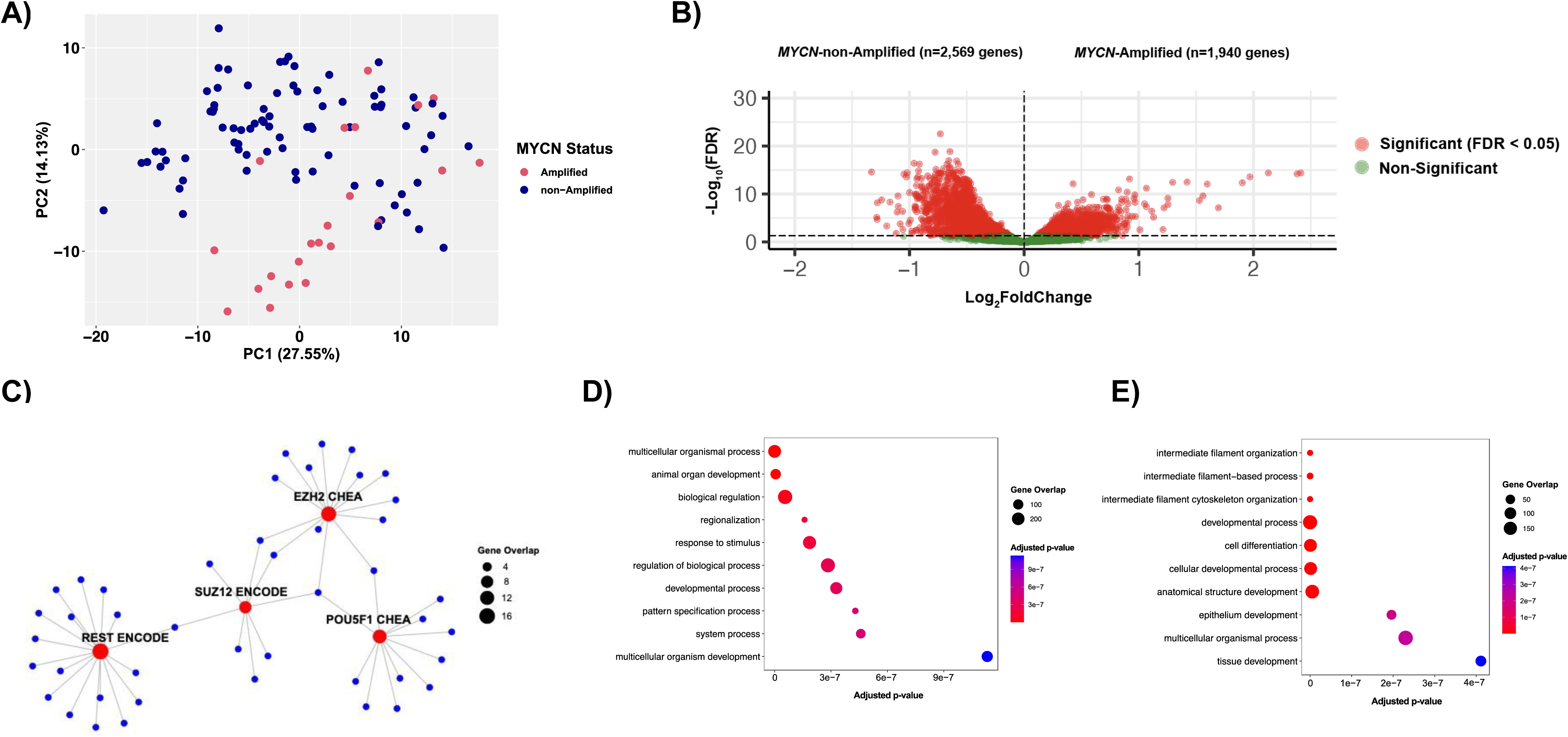
*MYCN*-amplified neuroblastoma tumors have increased 5-hmC on PRC2 target genes. **A)** Principal Component Analysis (PCA) of 5-hmC profiles in neuroblastoma patient derived *MYCN*-amplified tumors (n=22) and *MYCN*-non-amplified tumors (n=83). **B)** Differentially hydroxymethylated genes between *MYCN*-amplified and *MYCN*-non-amplified tumors. 1,940 genes had significantly higher (FDR<0.05) 5-hmC deposition in *MYCN*-amplified tumors, while 2,569 genes had significantly higher (FDR<0.05) 5-hmC in *MYCN*-non-amplified tumors. **C)** Transcription factor targets significantly enriched for in genes (n=451) with significantly increased 5-hmC deposition (in the top 10% by log_2_FoldChange (Log_2_FC)) in *MYCN*-amplified tumors compared to *MYCN-*non-amplified tumors, by gene set enrichment analysis (GSEA). Red nodes indicate significantly (FDR<0.05) enriched datasets, while blue nodes constitute the genes enriched for 5-hmC in *MYCN*-amplified tumors overlapping with the indicated publicly available datasets. **D)** Top 10 significant (FDR<0.05) Gene Ontology: Biological Processes (GO:BP) enriched for in genes (n=451) with significantly increased 5-hmC deposition (ie top 10% by log_2_FoldChange (Log_2_FC)) in *MYCN*-amplified tumors compared to *MYCN-*non-amplified tumors by GSEA. **E)** Top 10 significant (FDR<0.05) GO:BP enriched for in genes (n=450) with significantly increased 5-hmC deposition (top 10% by log_2_FoldChange (Log_2_FC)) in *MYCN*-non-amplified tumors compared to *MYCN-*amplified tumors by GSEA.

Genes (n=451) with significantly increased 5-hmC deposition in *MYCN*-amplified tumors were enriched for target genes of SUZ12, EZH2, REST, and POU5F1 (**FIGURE 1C**). In *MYCN*-amplified tumors, we also observed significant enrichment of 5-hmC on Gene Ontology Biological Processes pathways including “animal organ development,” “developmental process,” and “multicellular organism development”, consistent with prior studies (**FIGURE 1D**)^15,22^. Genes with significantly increased 5-hmC deposition in *MYCN*-non-amplified tumors (n=450) were not significantly enriched for any transcription factor pathways. Biological processes involving intermediate filament organization and to a lesser extent, development-related pathways, were enriched for 5-hmC in *MYCN*-non-amplified tumors (**FIGURE 1E**).

Because publicly available transcription factor datasets are built on several different model systems, including mESCs, we next investigated whether 5-hmC was enriched on neuroblastoma-specific PRC2 targets in *MYCN*-amplified tumors^23^. Genes with significantly increased 5-hmC deposition in *MYCN*-amplified tumors had significant overlap with a neuroblastoma specific H3K27me3-enriched gene set (p=0.003)^24^.

Conversely, genes with significantly increased 5-hmC deposition in *MYCN*-non-amplified tumors did not have significant overlap with H3K27me3-enriched genes (p=0.19). When we restricted our analysis to HR tumors, we again found that the genes with significantly increased 5-hmC deposition in *MYCN-*amplified HR tumors (n=21) had significant overlap with neuroblastoma specific H3K27me3-enriched genes (p<0.001), which was not the case in *MYCN*-non-amplified HR tumors (n=26) (p=0.53). Thus, we concluded that *MYCN*-amplified tumors had increased 5-hmC deposition on PRC2 target genes.

### Transcription start site 5-hmC and H3K27me3 directly co-localize at the nucleosomal level in MYCN-amplified neuroblastoma

To investigate the functional relationship between 5-hmC, PRC2, and *MYCN*-amplification, we analyzed genome-wide 5-hmC and H3K27me3 profiles in the *MYCN*-amplified SK-N-BE2, LA1-55n, LA1-5s, NBL-W-N, NBL-W-S; *MYCN* overexpressing NBL-S; the *MYC* overexpressing SH-SY5Y; and SHEP neuroblastoma cell lines. We found that cell lines with high *MYCN* or *MYC* expression had increased deposition of H3K27me3 across the transcription start sites (TSS) of protein-coding genes, and cell lines with low MYCN or MYC expression had relatively low levels of H3K27me3. While the extent of H3K27me3 deposition varied by *MYCN*/*MYC* expression, patterns of H3K27me3 deposition were relatively conserved across all eight included neuroblastoma cell lines (**FIGURE 2A**). Across all included cell lines, genes that were enriched for H3K27me3 had notable enrichment of 5-hmC at the TSS (**SUPPLEMENTAL FIGURE 2A-H**). Genes lacking H3K27me3 signal generally lacked 5-hmC at the TSS, with enrichment of 5-hmC noted 500-1000bp upstream and downstream of the TSS. Next, to determine the extent of overlap between H3K27me3 and 5-hmC across all included cell lines, H3K27me3 and 5-hmC peaks were called and assigned to genes if they were located within 3kb of TSSs. Through this analysis, we found that cell lines with the highest *MYCN*/*MYC* expression had significantly more 5-hmC/H3K27me3 co-enriched genes compared to cell lines with low *MYCN*/*MYC* expression (**FIGURE 2B**).

**FIGURE 2:**
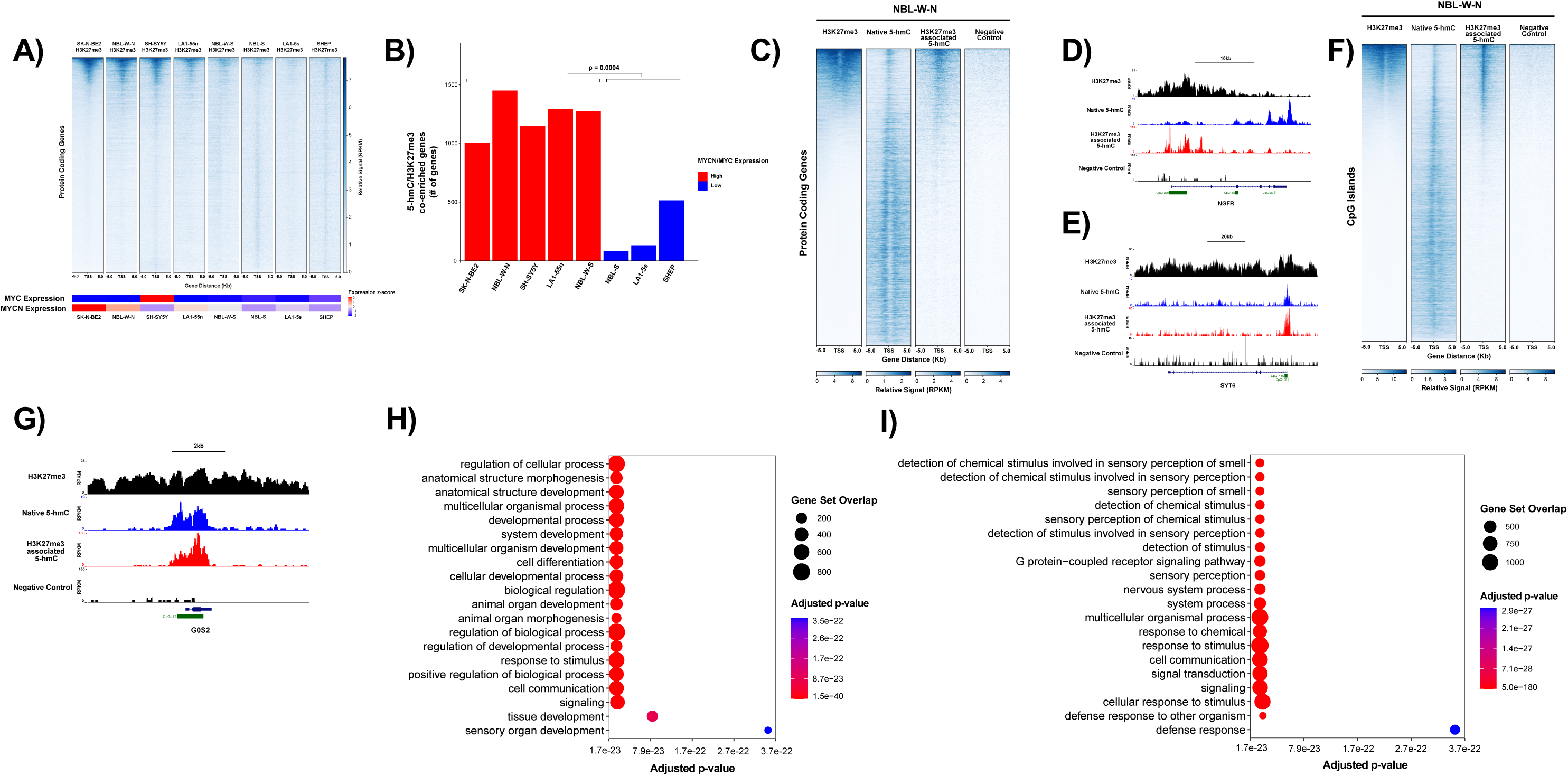
Transcription start site 5-hmC and H3K27me3 directly co-localize at the nucleosomal level in *MYCN*-amplified neuroblastoma. **A)** ChIP-Seq for H3K27me3 (normalized to reads per kilobase million (RPKM)) +/-5kb of the transcription start site (TSS) of protein coding genes, in the SK-N-BE2, NBL-W-N, SH-SY5Y, LA1-55n, NBL-W-S, NBL-S, LA1-5s, and SHEP cell lines. H3K27me3 profiles are ordered by signal in the SK-N-BE2 cell line. Below the H3K27me3 profiles, heatmaps are shown representing relative *MYCN* or *MYC* expression by z-score across included cell lines. **B)** Number of 5-hmC/H3K27me3 co-enriched genes across cell lines with high or low *MYCN*/*MYC* expression. Significance determined by two-tailed t-test. **C)** H3K27me3, native 5-hmC, H3K27me3 associated 5-hmC, and negative control profiles (normalized to RPKM) +/- 5kb of the TSS of protein coding genes in the NBL-W-N cell line. H3K27me3, native 5-hmC, H3K27me3 associated 5-hmC, and negative control tracks normalized to RPKM are shown for **D)** *NGFR* and **E)** *SYT6* in NBL-W-N. **F)** H3K27me3, native 5-hmC, H3K27me3 associated 5-hmC, and negative control profiles (normalized to RPKM) +/- 5kb of CpG islands in the NBL-W-N cell line. H3K27me3, native 5-hmC, H3K27me3 associated 5-hmC, and negative control tracks normalized to RPKM are shown for **G)** *G0S2* in NBL-W-N. **H)** Top 20 significant (FDR<0.05) GO:BP enriched for in genes (n=1,256) with 5-hmC/H3K27me3 co-enrichment in NBL-W-N. **I)** Top 20 significant (FDR < 0.05) GO:BP enriched for in genes (n=2,434) with H3K27me3-only enrichment in NBL-W-N.

Similar to the results we obtained, prior studies in mESCs have found associations of 5-hmC and H3K27me3 at TSSs of transcriptionally repressed genes through the analysis of bulk-sequencing experiments^18–20,25^. To determine whether 5-hmC and H3K27me3 could be deposited at the same genomic location and coexist on the same nucleosome in *MYCN*-amplified neuroblastoma, we performed serial CUT&RUN (C&R) for H3K27me3 followed by nano-hmC-Seal^26^ (hereafter referred to as H3K27me3-associated 5-hmC), enabling us to directly assess the relationship between 5-hmC and H3K27me3 at the nucleosomal level. C&R for H3K27me3 and hmC-Seal generated 5-hmC profiles (hereafter referred to as native 5-hmC) were utilized as references. In the *MYCN*-amplified NBL-W-N and SK-N-BE2 cell lines, H3K27me3-associated 5-hmC profiles closely matched the global profile of H3K27me3 at TSSs.

Both H3K27me3 and H3K27me3-associated 5-hmC had a relatively broad distribution over the TSSs of a subset of protein-coding genes, demonstrating that 5-hmC and H3K27me3 directly co-localize at the nucleosomal level (**FIGURE 2C**; **SUPPLEMENTAL FIGURE 3A**). A negative control in which DNA isolated from C&R for H3K27me3 passed through the typical nano-hmC-Seal workflow with the omission of the T4-Beta-Glucosyltransferase (T4-BGT) enzyme had negligible signal across protein-coding genes (**FIGURE 2C**; **SUPPLEMENTAL FIGURE 3A**). Interrogation of gene level coverage tracks normalized to reads per kilobase million (RPKM) further supported the direct association between 5-hmC and H3K27me3 at TSS of protein-coding genes, as demonstrated by *NGFR* (**FIGURE 2D**), a well-described EZH2 regulated tumor suppressor in neuroblastoma, and *SYT6* (**FIGURE 2E**), a gene belonging to the synaptotagmin family with low expression in neuroblastoma^27^. Interestingly, in both *NGFR* and *SYT6*, 5-hmC and H3K27me3 co-enrichment were noted at CpG island sites in the promoter region.

DNA methylation and H3K27me3 are mutually exclusive at CpG island sites in mESCs^28^. Given the co-localization of 5-hmC and H3K27me3 at CpG islands of two candidate genes, we next formally evaluated the presence of 5-hmC and H3K27me3 at CpG islands. As expected, we found that a subset of CpG islands contained enrichment of H3K27me3 in both NBL-W-N (**FIGURE 2F**) and SK-N-BE2 (**SUPPLEMENTAL FIGURE 3B**). In both cell lines, H3K27me3-associated 5-hmC signal strongly overlapped with that of H3K27me3 at CpG island sites, suggesting that CpG islands marked by H3K27me3 were selectively enriched for 5-hmC. The direct association between 5-hmC and H3K27me3 at CpG island sites was further confirmed through interrogation of gene level coverage tracks (**FIGURE 2G**).

We investigated the biological significance of 5-hmC/H3K27me3 co-enriched genes. To be defined as a gene with 5-hmC/H3K27me3 enrichment, a gene had to be enriched for H3K27me3, native 5-hmC, and H3K27me3 associated 5-hmC. We identified 1,256 genes co-enriched for 5-hmC/H3K27me3 while 2,434 genes were enriched for H3K27me3-only in the NBL-W-N cell line (**SUPPLEMENTAL FIGURE 4A**). In SK-N-BE2, we identified 1,937 genes co-enriched for 5-hmC/H3K27me3 and 2,169 genes enriched for H3K27me3 (**SUPPLEMENTAL FIGURE 3C**). We found that 5-hmC/H3K27me3 co-enriched genes were significantly enriched for biological pathways including “anatomical structure development,” “developmental process,” and “cell differentiation” in both NBL-W-N (**FIGURE 2H**) and SK-N-BE2 (**SUPPLEMENTAL FIGURE 3D**). In comparison, H3K27me3-only genes were enriched for pathways such as “sensory perception” and “G protein-coupled receptor signaling pathways” in both NBL-W-N (**FIGURE 2I**) and SK-N-BE2 (**SUPPLEMENTAL FIGURE 3E**). Using RNA-Seq of NBL-W-N, we found that both 5-hmC/H3K27me3 co-enriched genes and H3K27me3-only genes were transcriptionally repressed (mean log_2_(TPM+1) of 1.19 and 0.80, respectively) in comparison to highly expressed genes (mean log_2_(TPM + 1) of 7.93) (**SUPPLEMENTAL FIGURE 4B**). Similar results were obtained in the SK-N-BE2 cell line, where both 5-hmC/H3K27me3 co-enriched genes and H3K27me3-only genes were also transcriptionally repressed (mean log_2_(TPM+1) of 1.39 and 0.92, respectively) in comparison to highly expressed genes (mean log_2_(TPM + 1) of 7.95) (**SUPPLEMENTAL FIGURE 3F**).

### Pharmacologic inhibition of PRC2 reduces H3K27me3 but does not alter 5-hmC deposition

Having established the nucleosomal level co-localization between 5-hmC and H3K27me3 and the biological relevance of genes co-enriched for 5-hmC/H3K27me3, we next probed the functional relationship between these two marks in *MYCN*-amplified neuroblastoma. First, we investigated whether modulation of H3K27me3 deposition through PRC2 inhibition with tazemetostat, an FDA-approved EZH2 inhibitor, resulted in alteration of 5-hmC deposition in NBL-W-N and SK-N-BE2. As expected, tazemetostat treatment resulted in a notable reduction in levels of H3K27me3 in both NBL-W-N and SK-N-BE2 (**FIGURE 3A, B**). Through RNA-Seq, we found that following tazemetostat treatment, 1,164 and 318 genes were significantly upregulated (FDR < 0.05), and 464 and 18 genes were significantly downregulated (FDR < 0.05) in NBL-W-N and SK-N-BE2, respectively (**FIGURE 3C**). Consistent with prior reports, pathways such as “nervous system development” and “neurogenesis” were significantly upregulated following treatment with tazemetostat in both NBL-W-N and SK-N-BE2 (**FIGURE** 3D)^10,29,30^.

**FIGURE 3:**
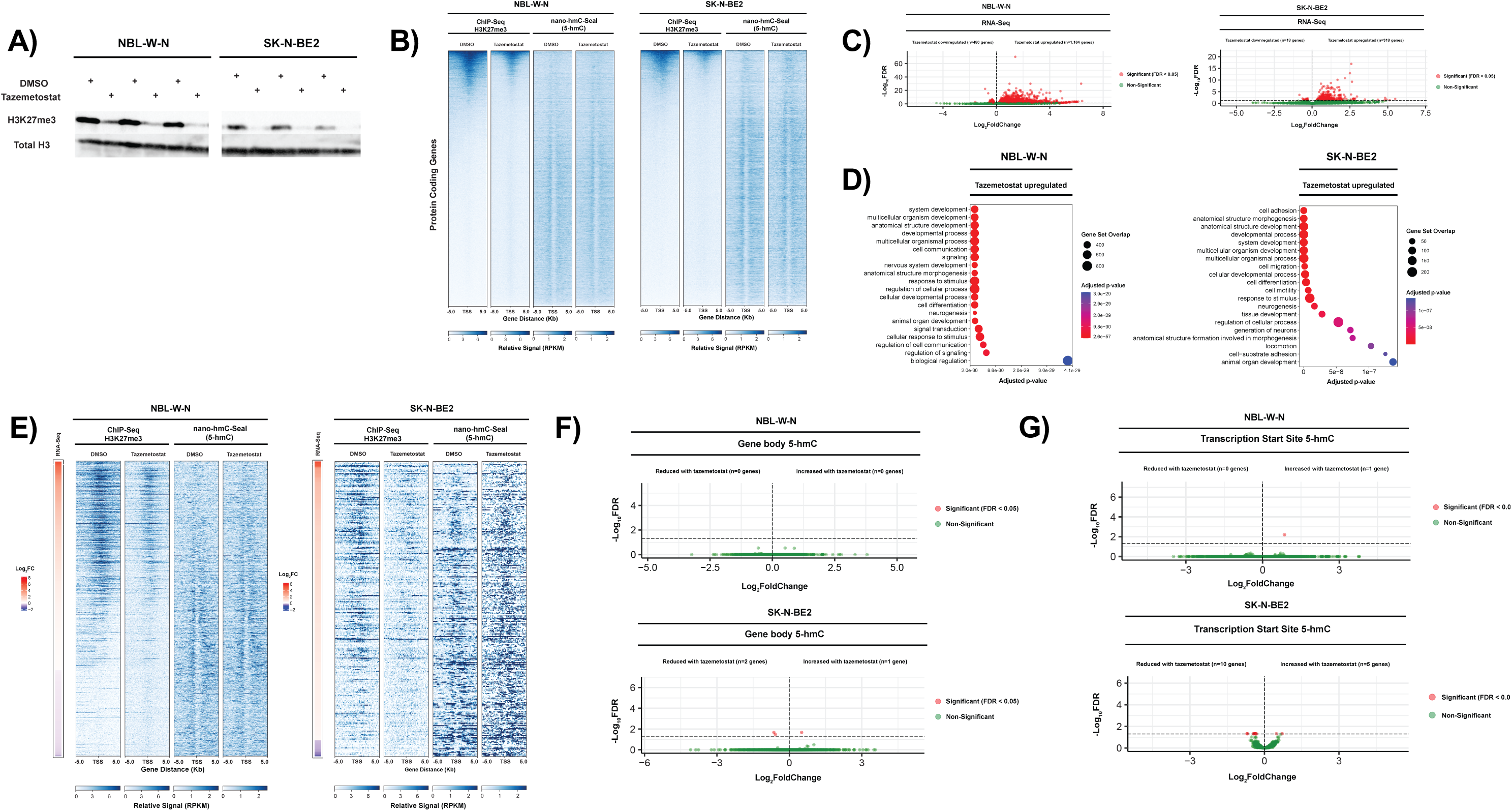
Pharmacologic inhibition of PRC2 reduces H3K27me3 but does not alter 5-hmC deposition in NBL-W-N and SK-N-BE2. **A)** Western blot showing relative levels of H3K27me3 and total H3 following treatment with tazemetostat in NBL-W-N (left panel) and SK-N-BE2 (right panel). **B)** H3K27me3 and 5-hmC profiles of protein coding genes in DMSO and tazemetostat treated NBL-W-N (left panel) and SK-N-BE2 (right panel). For both NBL-W-N and SK-N-BE2, samples are ranked according to the relative signal in the respective DMSO treated ChIP-Seq H3K27me3 profiles. **C)** Differentially expressed genes by RNA-Seq following treatment with tazemetostat in NBL-W-N (left panel) and SK-N-BE2 (right panel). **D)** Top 20 significant (FDR<0.05) GO:BP enriched in genes with significantly increased expression following treatment with tazemetostat in NBL-W-N (left panel) and SK-N-BE2 (right panel). **E)** H3K27me3 and 5-hmC profiles of genes with significant changes in expression following treatment with tazemetostat in NBL-W-N (left panel) and SK-N-BE2 (right panel). Genes are presented in decreasing order by log_2_FoldChange in expression following treatment with tazemetostat. Heatmap to the left of the figure represents log_2_FoldChange in expression following treatment with tazemetostat. **F)** Differential hydroxymethylation across the gene body following treatment with tazemetostat in NBL-W-N (top panel) and SK-N-BE2 (bottom panel). **G)** Differential hydroxymethylation within 3kb of the TSS following treatment with tazemetostat in NBL-W-N (top panel) and SK-N-BE2 (bottom panel).

Next, we investigated changes in 5-hmC through nano-hmC-Seal following tazemetostat treatment. Though tazemetostat notably decreased levels of H3K27me3, there were no global changes in 5-hmC deposition in both NBL-W-N and SK-N-BE2 (**FIGURE 3B**). Further, when examining the subset of genes with transcriptional change in NBL-W-N (n=1,628 genes) and SK-N-BE2 (n=336 genes) following tazemetostat treatment, we noted the expected reduction in H3K27me3 without any apparent changes in 5-hmC deposition (**FIGURE 3E**). To formally investigate changes in 5-hmC following PRC2 inhibition with tazemetostat, 5-hmC reads were counted across gene bodies and +/- 3kb of the TSS of protein coding genes. In both NBL-W-N and SK-N-BE2, changes in 5-hmC were insufficient to explain the majority of transcriptional change induced by PRC2 pharmacologic inhibition (**FIGURE 3F, G**). Therefore, we concluded that pharmacologic inhibition of PRC2 did not alter patterns of 5-hmC deposition in *MYCN*-amplified neuroblastoma cell lines.

### Chemical inhibition of 5-hmC deposition with cobalt chloride results in a loss of H3K27me3 over protein-coding genes in neuroblastoma

Subsequently, we investigated whether modulation of 5-hmC deposition would alter patterns of H3K27me3 deposition in *MYCN-*amplified neuroblastoma using the SK-N-BE2 cell line. Given that the TET proteins belong to the class of Fe(II) and 2-oxoglutarate (2-OG) dependent dioxygenases, we posited that treatment of cells with cobalt chloride would chemically inhibit the activity of TET enzymes resulting in a global loss of 5-hmC^31–33^. Indeed, treatment of the SK-N-BE2 cell line with cobalt chloride diminished global levels of 5-hmC in a dose-dependent manner (**FIGURE 4A**). Treatment of SK-N-BE2 with cobalt chloride did not affect the expression of EZH2, EED, or SUZ12, and did not result in decreased global levels of H3K27me3 (**FIGURE 4B**).

**FIGURE 4:**
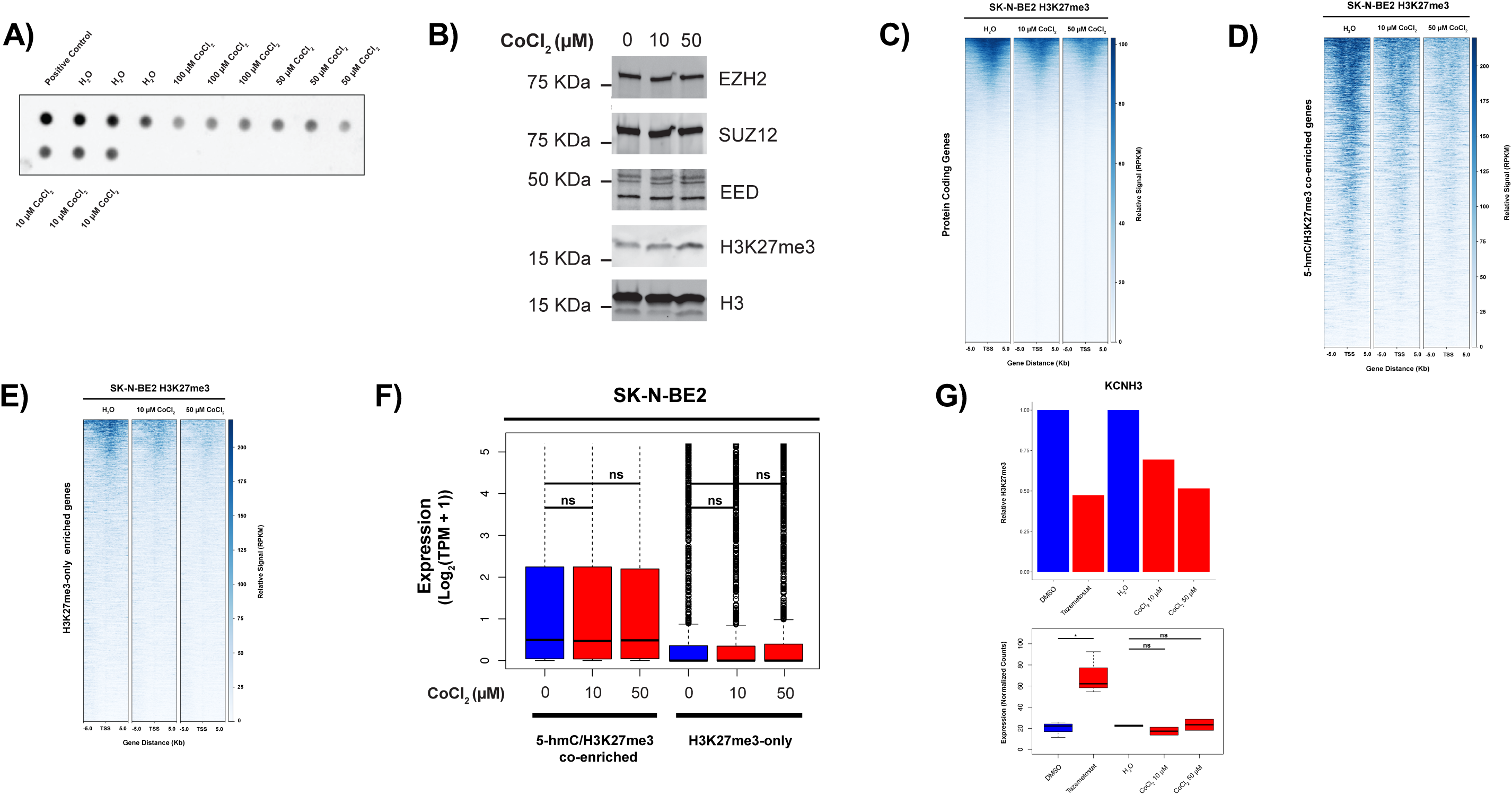
Chemical inhibition of 5-hmC deposition with cobalt chloride results in a loss of H3K27me3 over protein-coding genes in SK-N-BE2. **A)** Dot blot showing relative levels of 5-hmC following treatment with cobalt chloride in SK-N-BE2. Mouse embryonic stem cell DNA was utilized as a positive control. **B)** Western blot showing levels of EZH2, SUZ12, EED, H3K27me3 and total H3 following treatment with cobalt chloride in SK-N-BE2. H3K27me3 of **C)** protein coding genes, **D)** 5-hmC/H3K27me3 co-enriched genes, and **E)** H3K27me3-only enriched genes in H_2_O and cobalt chloride treated SK-N-BE2. Genes are ordered according to the H_2_O CUT&RUN H3K27me3 profile. **F)** Expression across 5-hmC/H3K27me3 co-enriched and H3K27me3-only enriched genes following treatment with cobalt chloride in SK-N-BE2. No within group cobalt chloride versus H_2_O comparisons were significant. ‘ns’ denotes not significant. **G)** Relative deposition of H3K27me3 across *KCNH3* (top) and expression of *KCNH3* (bottom) in DMSO, tazemetostat, H_2_O, and CoCl_2_ treated SK-N-BE2.

We next determined locus-specific changes in H3K27me3 deposition and accompanying transcriptional changes following loss of 5-hmC with cobalt chloride treatment. Treatment with cobalt chloride resulted in a loss of H3K27me3 over protein-coding genes, 5-hmC/H3K27me3 co-enriched genes, and H3K27me3-only enriched genes (**FIGURE 4C, D, E**). Despite this, there was no significant change in the expression of 5-hmC/H3K27me3 enriched genes or H3K27me3-only enriched genes following treatment with cobalt chloride, suggesting that alternative mechanisms maintain the transcriptional repression of these genes in the absence of H3K27me3 (**FIGURE 4F**). Interrogation of candidate genes further demonstrate this phenomenon, whereby loss H3K27me3 following treatment with cobalt chloride did not result in transcriptional activation of H3K27me3 targets despite having transcriptional activation with loss of H3K27me3 following treatment with tazemetostat (**FIGURE 4G, SUPPLEMENTAL FIGURE 5A-I**).

### Genetic inactivation of *TET2* reduces H3K27me3 signal at 5-hmC-H3K27me3 positive promoters and increases DNA methylation levels

To further investigate the specific role that 5-hmC directly played in the deposition of H3K27me3, we genetically inactivated the *TET* genes in the SK-N-BE2 neuroblastoma cell line. We first proceeded with a double knockout of the *TET1* and *TET3* genes because of their higher transcript levels in these cells (data not shown). Combined genetic inactivation of *TET1* and *TET3* did not result in any significant change in the global levels of 5-hmC compared to control gRNA (**SUPPLEMENTAL FIGURE 6A-B**). At this point we genetically inactivated *TET2*, the only other known enzyme that can deposit 5-hmC in mammalian cells, in combination with *TET3*^34^.

Though we utilized CRISPR/Cas9 with sgRNAs against *TET2* and *TET3*, only TET2 was decreased at the protein level, thus creating a cell line we hereafter refer to as *TET2* knockout (KO) cells (**FIGURE 5A, SUPPLEMENTAL FIGURE 7**). *TET2* KO cells had substantially decreased 5-hmC levels compared to controls (**FIGURE 5B**). To understand whether this global loss of 5-hmC affected PRC2 mediated deposition of H3K27me3, we performed C&R for H3K27me3 in *TET2* KO cells. We found a drastic reduction of the levels of the H3K27me3 deposition in *TET2* KO cells compared to controls at many TSS (**FIGURE 5C**). Most importantly, loss of H3K27me3 was more profound around 5-hmC/H3K27me3 positive regions in comparison to H3K27me3-only regions (**FIGURE 5C** and **SUPPLEMENTAL FIGURE 8**). As expected, the minor reduction of the H3K27me3 levels at H3K27me3-only genes did not elicit a significant transcriptional response (**FIGURE 5D**). Interestingly, 5-hmC/H3K27me3 positive genes also did not show any significant changes in their transcription levels despite a drastic (∼50%) reduction in H3K27me3 signal, consistent with our cobalt chloride experiments (**FIGURE 5D**).

**FIGURE 5:**
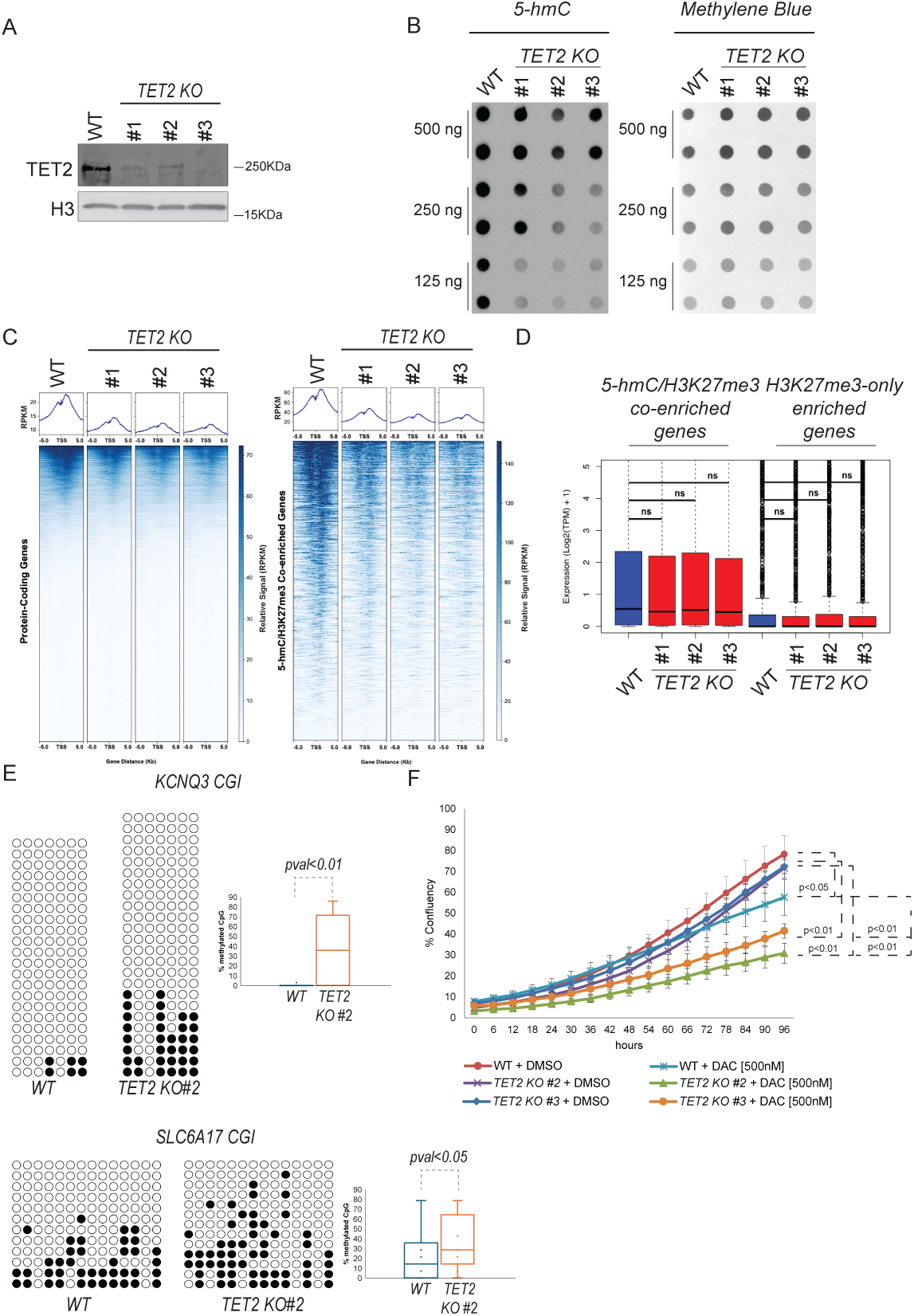
Genetic inactivation of *TET2* reduces H3K27me3 signal at 5-hmC-H3K27me3 positive promoters and increases DNA methylation levels. **A)** Western blot showing relative expression of TET2 in SK-N-BE2 wild-type (WT) and SK-N-BE2 *TET2* knockout (KO) clones (#1, 2, 3). **B)** Dot-blot showing relative 5-hmC (left) and total DNA by methylene blue stain (right) in SK-N-BE2 WT and SK-N-BE2 *TET2* KO cells. Loading amounts of DNA as indicated by the panel on the left. **C)** H3K27me3 of protein coding genes and 5-hmC/H3k27me3 co-enriched genes in WT and *TET2* KO SK-N-BE2 cells. **D)** Expression across 5-hmC/H3K27me3 co-enriched and H3K27me3-only enriched genes in WT and *TET2* KO SK-N-BE2 cells. ‘ns’ denotes not significant. **E)** Bisulfite PCR at CpG islands near the TSS on *KCNQ3* and *SLC6A17* in WT and *TET2* KO SK-N-BE2. Filled in circles represented methylated CpG while unfilled circles represent unmethylated CpG. **F)** Percentage confluency in WT and *TET2* KO SK-N-BE2 cells treated with DMSO or decitabine 500nM.

We hypothesized that an increase in 5-mC following *TET2* KO may be a plausible explanation for the maintenance of transcriptional repression at 5-hmC/H3K27me3 sites. To test this, we chose two genes marked at the TSS by concomitant 5-hmC/H3K27me3: *KCNQ3* and *SLC6A17*. Bisulfite PCR at CpG islands near the TSS confirmed an increase in 5-mC at those loci in *TET2* KO cells compared to controls (**FIGURE 5E**). We also determined that *TET2 KO* cells not only have increased focal 5-mC, but they also become more sensitive than control cells to the DNA hypomethylating agent decitabine (**FIGURE 5F**). Altogether these data suggest that TET2 deposited 5-hmC is important for H3K27me3 deposition at 5-hmC/H3K27me3 regions, and its depletion leads to 5-mC accumulation, making cells more sensitive to decitabine.

### 5-hmC deposition predisposes H3K27me3 enriched genes to transcriptional activation following PRC2 inhibition

To further characterize how 5-hmC deposition alters the transcriptional behavior of H3K27me3-marked genes, we examined transcriptional changes of the previously identified 5-hmC/H3K27me3 co-enriched (n=1,256; n=1,937) and H3K27me3-only enriched (n=2,434; n=2,169) genes in the NBL-W-N and SK-N-BE2 cell lines respectively following PRC2 inhibition with tazemetostat. Both 5-hmC/H3K27me3 and H3K27me3-only genes had notable reductions in H3K27me3 following treatment with tazemetostat (**FIGURE 6A, B**). 5-hmC/H3K27me3 co-enriched genes had significant overlap with genes upregulated by tazemetostat in both NBL-W-N and SK-N-BE2 (p=1.5e-108; p=2.8e-43). In contrast, no significant overlap was detected between H3K27me3-only enriched genes and genes upregulated by tazemetostat in both NBL-W-N and SK-N-BE2 (p=0.89 and p=0.71). Furthermore, 5-hmC/H3K27me3 co-enriched genes had significantly increased expression (p=0.0004 and p=0.007) while PRC2-only enriched genes did not significantly change in expression (p=0.21 and p=0.45) following treatment with tazemetostat respectively in NBL-W-N and SK-N-BE2 (**FIGURE 6C**). Thus, we concluded that 5-hmC deposition at the TSS predisposes H3K27me3 marked genes to transcriptional activation upon PRC2 inhibition with tazemetostat.

**FIGURE 6:**
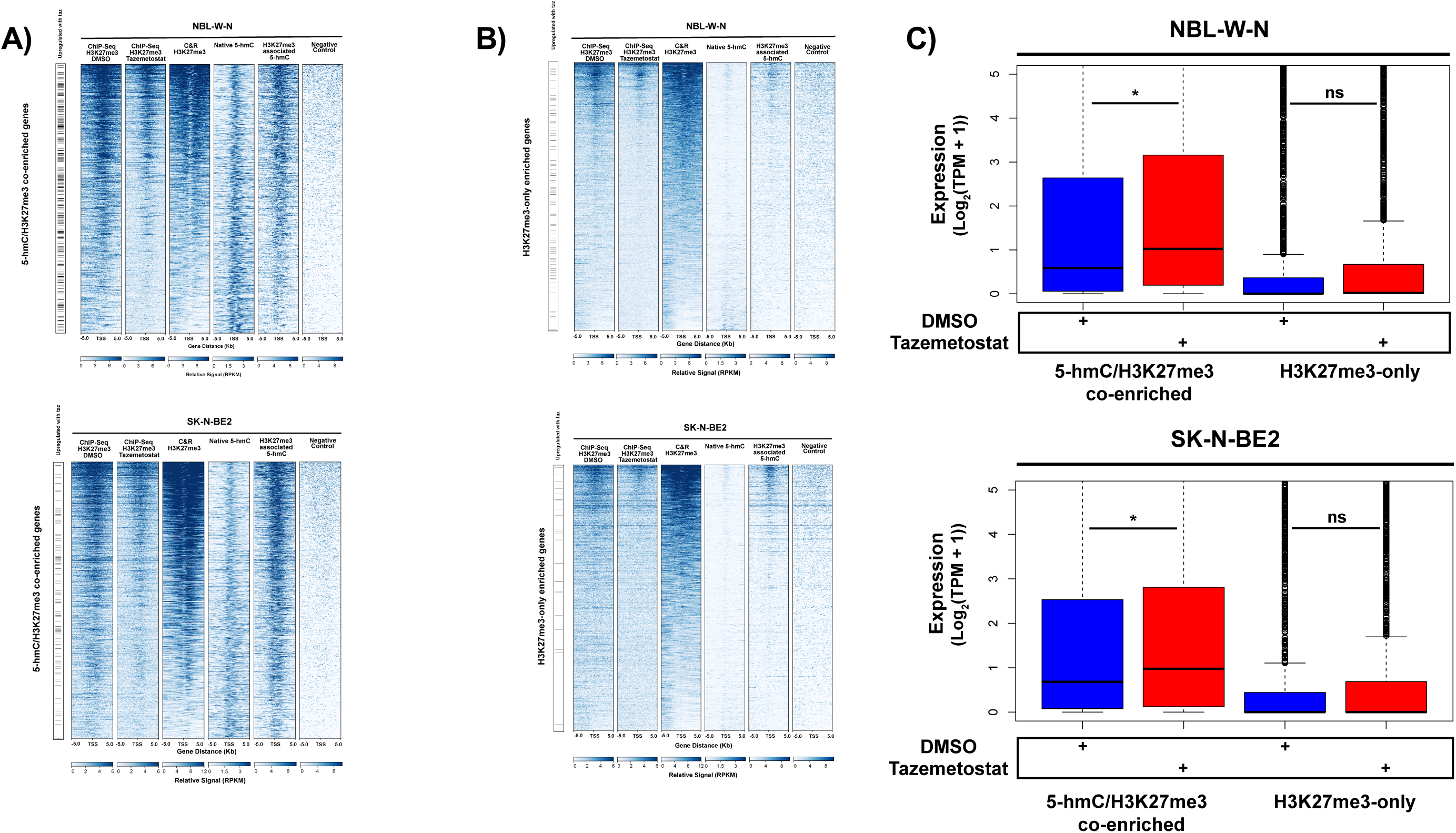
5-hmC deposition predisposes H3K27me3 enriched genes to transcriptional activation following PRC2 inhibition. **A)** DMSO treated ChIP-Seq H3K27me3, tazemetostat treated ChIP-Seq H3K27me3, Cut&Run (C&R) H3K27me3, native 5-hmC, H3K27me3 associated 5-hmC, and negative control profiles of 5-hmC/H3K27me3 co-enriched genes, ranked by C&R H3K27me3, in NBL-W-N (n=1,256 genes) and SK-N-BE2 (1,937 genes). Black dashes to the left of each figure indicate genes that significantly gained expression following treatment with tazemetostat in NBL-W-N (n=294 genes) and SK-N-BE2 (n=121 genes). **B)** H3K27me3, native 5-hmC, H3K27me3 associated 5-hmC, and negative control profiles of H3K27me3-only enriched genes, ranked by H3K27me3, in NBL-W-N (n=2,434 genes) and SK-N-BE2 (2,169 genes). Black dashes to the left of the figure indicate genes (n=126 genes) that significantly gained expression following treatment with tazemetostat in NBL-W-N and SK-N-BE2 (n=31 genes). **C)** Expression across 5-hmC/H3K27me3 enriched and H3K27me3-only enriched genes following treatment with tazemetostat in NBL-W-N and SK-N-BE2. * denotes p-value < 0.05, and ns denotes not significant. P-value determined through two-tailed t-test.

### Low expression of 5-hmC/H3K27me3 co-enriched genes is associated with poor clinical outcome in patients with neuroblastoma

Finally, we investigated the clinical significance of genes co-enriched for 5-hmC and H3K27me3 in two distinct cohorts of neuroblastoma patients^35,36^. A consensus set of n=733 genes with 5-hmC/H3K27me3 co-enrichment in SK-N-BE2 and NBL-W-N was identified. Gene set variance analysis (GSVA) was utilized to generate 5-hmC/H3K27me3 signature scores, with lower signature scores indicating relatively low expression of 5-hmC/H3K27me3 co-enriched genes^37^. GSVA was also utilized to generate MYCN signature scores using a previously described gene signature^38^.

In the SEQC-NB dataset (n = 493), MYCN signature scores were significantly negatively correlated with 5-hmC/H3K27me3 signature scores (R=-0.6, p<2.2e^-16^) (**FIGURE 7A**)^36^. Further, INSS Stage 4 tumors (n=181) had significantly lower 5-hmC/H3K27me3 signatures scores compared to Stage 1, 2, 3, and 4S tumors (n=312; p=3.4e^-8^) (**FIGURE 7B**). To assess the relationship between 5-hmC/H3K27me3 signature and overall survival (OS), a 5-hmC/H3K27me3 signature score of -0.15 was utilized to classify samples as “5-hmC/H3K27me3 low expression” or “5-hmC/H3K27me3 high expression.” In the full SEQC-NB cohort, samples from patients with low expression of 5-hmC/H3K27me3 co-enriched genes had significantly worse OS compared to samples from patients with high expression of 5-hmC/H3K27me3 co-enriched genes (5-year OS (60.7% [95% CI: 52.5%-70.1%] vs 85.1% [95% CI: 81.3%-89.1%]; p < 0.0001)) (**FIGURE 7C**). In the subset of patients with HR disease (n=175), samples from patients with low expression of 5-hmC/H3K27me3 co-enriched genes also had significantly worse OS compared to samples from patients with high expression of 5-hmC/H3K27me3 co-enriched genes (5-year OS (34.6% [95% CI: 24.9%-48.2%] vs 51.4% [95% CI: 41.3%-63.9%]; p=0.0028)) (**FIGURE 7D**).

**FIGURE 7:**
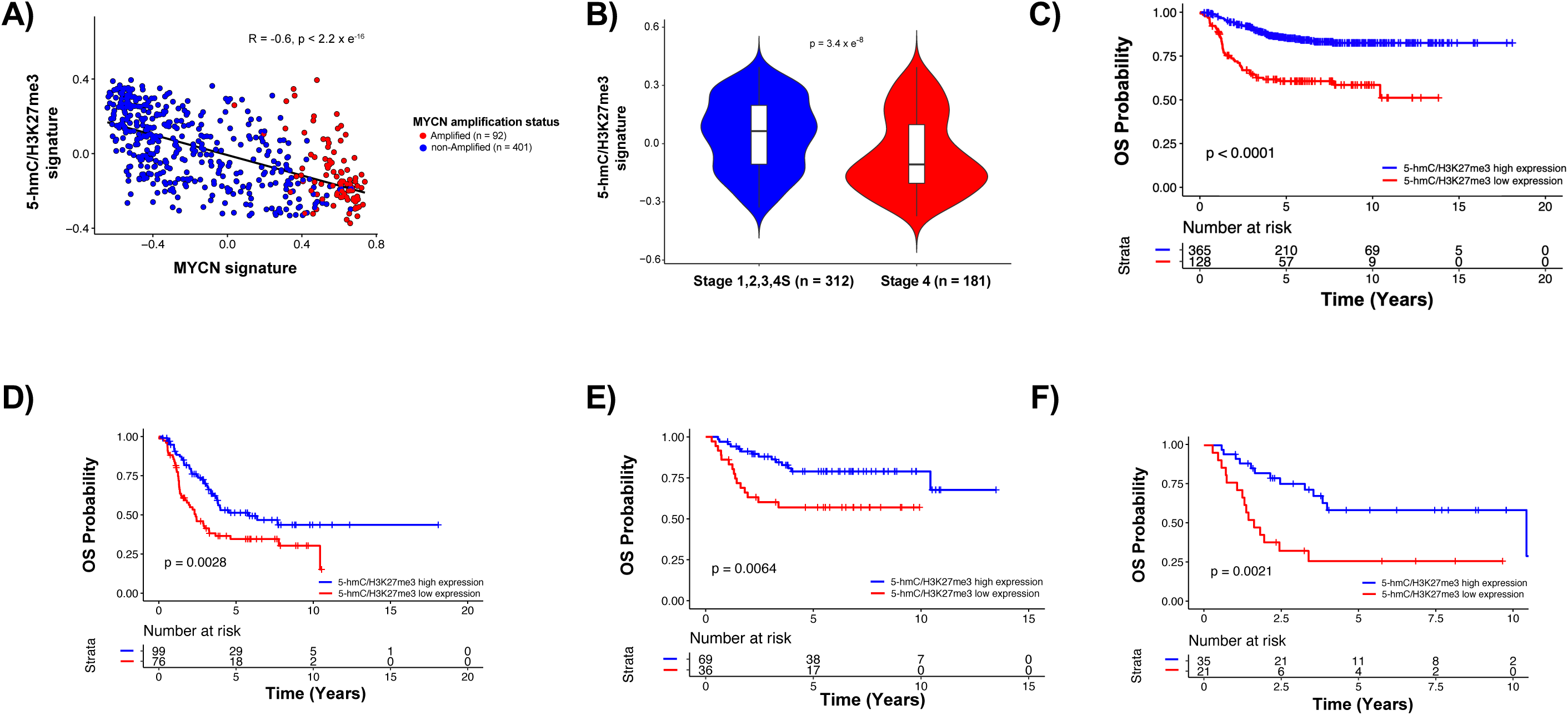
Low expression of 5-hmC/co-enriched genes is associated with poor clinical outcome in patients with neuroblastoma. Expression and phenotypic data for 493 patients (SEQC-NB cohort) and 105 patients (GSE73517) were obtained from R2.amc.nl. **A)** Scatter plot of 5-hmC/H3K27me3 signature against MYCN signature in the SEQC-NB cohort. **B)** 5-hmC/H3K27me3 signature by INSS Stage in patients with neuroblastoma in the SEQC-NB cohort. Significance determined by two-tailed t-test. **C)** Kaplan-Meier curves depicting overall survival (OS) in all 493 patients in the SEQC-NB cohort by relative expression of 5-hmC/H3K27me3 co-enriched genes. **D)** Kaplan-Meier curves depicting OS in 175 patients with high-risk disease in the SEQC-NB cohort by relative expression of 5-hmC/H3K27me3 co-enriched genes. **E)** Kaplan-Meier curves depicting OS in all 105 patients in the GSE73517 cohort by relative expression of 5-hmC/H3K27me3 co-enriched genes. **F)** Kaplan-Meier curves depicting OS in 56 patients with high-risk disease in the GSE73517cohort by relative expression of 5-hmC/H3K27me3 co-enriched genes. Significance of Kaplan-Meier curves determined by log-rank test.

Similar results were observed in a distinct cohort of 105 neuroblastoma tumors (GSE73518)^35^. Using the same cutoff score utilized in the SEQC-NB dataset, samples were classified as “5-hmC/H3K27me3 low expression” or “5-hmC/H3K27me3 high expression.” In the full GSE73518 cohort, samples from patients with low expression of 5-hmC/H3K27me3 co-enriched genes had significantly worse OS compared to samples from patients with high expression of 5-hmC/H3K27me3 co-enriched genes (5-year OS (57.0% [95% CI: 42.6%-76.2%] vs 78.9% [95% CI: 69.3%-89.9%]; p=0.0064)) (**FIGURE 7E**). In the subset of patients with HR disease (n=56), samples from patients with low expression of 5-hmC/H3K27me3 co-enriched genes also had significantly worse OS compared to samples from patients with high expression of 5-hmC/H3K27me3 co-enriched genes (5-year OS (26.1% [95% CI: 12.2%-56.0%] vs 58.7% [95% CI: 42.7%-80.6%]; p=0.0021)) (**FIGURE 7F**). Altogether, these results suggest that co-occupancy of 5-hmC/H3K27me3 on the gene set identified in the SK-N-BE2 and NBL-W-N cell lines is robustly associated with worse clinical outcome in patients with neuroblastoma.

## DISCUSSION

In the current study, we demonstrate that 5-hmC and H3K27me3, the catalytic product of PRC2, directly co-localize at the nucleosomal level of transcriptionally repressed regulators of development in *MYCN*-amplified neuroblastoma^15,39^. Chemical inhibition of 5-hmC deposition with cobalt chloride resulted in a loss of H3K27me3 over protein-coding genes. Genetic inhibition of 5-hmC deposition with *TET2* KO resulted in a loss of H3K27me3 at 5-hmC/H3K27me3 co-enriched genes, increased 5-mC accumulation, and increased sensitivity to the DNA hypomethylating agent decitabine. Furthermore, genes with 5-hmC/H3K27me3 co-localization were selectively vulnerable to transcriptional activation following treatment with tazemetostat, and low expression of 5-hmC/H3K27me3 co-enriched genes was associated with worse clinical outcome in two distinct cohorts of neuroblastoma patients.

Our results add to a growing body of literature exploring the relationship between DNA modifications and chromatin modifications. Through serial C&R for H3K27me3 followed by nano-hmC-Seal, we showed, for the first time, that 5-hmC and H3K27me3 directly co-localize at the nucleosomal level across a subset of CpG islands and TSSs of transcriptionally repressed regulators of development in *MYCN*-amplified neuroblastoma. Prior studies utilized ChIP-BS-seq to demonstrate a mutual antagonism between 5-mC and H3K27me3 at CpG island sites and promoter regions in mESCs^28,40,41^. Given that bisulfite sequencing is unable to distinguish between 5-mC and 5-hmC, these prior studies describe a potential mutual antagonism between both 5-mC/5-hmC and H3K27me3 in mESCs. However, in the current study, we found that 5-hmC and H3K27me3 are compatible in the context of both CpG islands and TSSs of a subset of protein coding genes. Our approach is advantageous in that we avoid cross-linking of DNA and are thus able to demonstrate with high specificity the direct nucleosomal co-localization of 5-hmC and H3K27me3^28,42^. Additionally, it was recently shown that in the context of trophoblast stem cells (TSCs), 5-mC/5-hmC and H3K27me3 can co-exist at CpG islands in contrast to the mutual exclusivity previously described in mESCs^42^. Thus, it is possible that the co-existence of 5-hmC and H3K27me3 depends on cell context, and that in more differentiated, yet still stem-like, cell states compared to mESCs, the two marks can directly co-localize at CpG islands.

We showed that inhibition of 5-hmC deposition with either cobalt chloride or *TET2* KO results in a loss of H3K27me3 over protein coding genes, suggesting that mechanistically, 5-hmC acts upstream of PRC2 to regulate transcriptional repression in *MYCN*-amplified neuroblastoma. This model is further supported by a recent study which demonstrated that site-directed demethylation with a dCas9-SunTag/scFv-TET1 catalytic domain construct resulted in a site-specific increase in H3K27me3 in mESCs, though it remains to be demonstrated whether this effect was mediated by 5-hmC^43^.

This mechanistic link between 5-hmC and PRC2 has possible therapeutic implications. First, we showed that 5-hmC/H3K27me3 co-enriched genes are selectively vulnerable to transcriptional activation following treatment with tazemetostat, highlighting a previously uncharacterized mechanism underlying the potential therapeutic benefit of EZH2 inhibition in neuroblastoma^10,29,30,44^. Additionally, a recent study showed that in adult T-cell leukemia/lymphoma, at least one patient developed resistance to valemetostat, a novel EZH1-EZH2 inhibitor, through the acquisition of a *TET2* loss of function mutation and subsequent accumulation of DNA methylation resulting in transcriptional repression^45^. In the current study, we showed that *TET2* KO cells had increased sensitivity to the DNA hypomethylating agent decitabine, thus demonstrating a possible strategy to bypass acquired resistance to EZH2 inhibition that may be applicable to contexts beyond neuroblastoma.

Given our results, we speculate that in neuronal differentiation during the process of DNA demethylation, PRC2 could be aberrantly recruited at 5-hmC occupied CpG islands (intermediate state of demethylation) therefore aberrantly silencing genes that were in the process of being expressed (subsequent to full DNA demethylation)^46^. This hypothesis is supported by three key pieces of information. First, early overexpression of MYCN during development results in the acquisition of embryonal stem-like qualities^22^. Recent studies have also highlighted the transcriptomic similarities between *MYCN-*amplified neuroblastoma and early neuroblasts/bridge cells found during development, in contrast to LR neuroblastoma which more closely resemble committed neuroblasts found at later stages of differentiation^47,48^. These studies support the idea that *MYCN-*amplified neuroblastoma is driven by errors occurring at early stages in development. Second, we found that 5-hmC/H3K27me3 co-enriched genes were preferentially expressed in comparison to H3K27me3-only enriched genes following PRC2 inhibition with tazemetostat. This suggests that the transcriptional machinery to activate 5-hmC/H3K27me3 co-enriched genes is already in place, underscoring the possibility that these genes were aberrantly silenced during the process of differentiation. Third, in neuroblastoma clinical cohorts, we found a strong negative correlation between *MYCN*-amplification and expression of 5-hmC/H3K27me3 co-enriched genes, further supporting the notion that 5-hmC/H3K27me3 co-enriched genes may have been aberrantly silenced during differentiation.

In summary, our results contribute to a growing understanding of the epigenomic underpinnings of *MYCN*-amplified neuroblastoma. We show that 5-hmC and H3K27me3 co-operate to repress mediators of development, highlighting a novel link between DNA modifications and chromatin modifications with potential therapeutic implications in *MYCN*-amplified neuroblastoma.

## METHODS

### Tumor biopsy derived genome-wide 5-hmC profile analysis

Raw phenotypic information and 5-hmC sequencing data for neuroblastoma tumor samples (n=107) included in the current study were downloaded from dbGaP accession number: phs001831.v1.p1. Fastqc (version 0.11.5) with default settings was used to assess raw read quality^49^. Raw reads with a minimum Phred score of 33 were filtered with Trimmomatic (version 0.36)^50^. Reads with leading and trailing bases below quality 3 and less than 36 base pairs in length were also trimmed. Bowtie2 (version 2.3.0) with default settings was used to align reads to GRCh38^51^. Picard (version 2.18.29) with default settings was utilized to remove duplicate reads^52^. Finally, featureCounts of subread (version 1.5.3) with the gene flag was used to count aligned reads across an entire gene body^53^. ggfortify (version 0.4.14) was utilized to conduct Principal Component Analysis (PCA)^54^.

DESeq2 (version 1.28.1) was utilized to identify genes with differential 5-hmC across included samples^55^. During differential 5-hmC analysis, age and sex were included as covariates. A false discovery rate (FDR) of less than 0.05 was considered significant. Volcano plots were generated with *EnhancedVolcano* (version 1.10.0)^56^.

ComplexHeatMap package (version 2.4.3) was utilized to perform unsupervised hierarchical clustering of tumor 5-hmC profiles (n=107 samples)^57^. Gene set enrichment analysis of genes in the top 10% by log-fold-change of significantly differentially hydroxymethyalted genes between *MYCN* amplified and non-amplified tumors was performed with ENRICHR, g:Profiler, and clusterProfiler^58–60^.

### Cell Culture, Tazemetostat, cobalt chloride, and decitabine treatment

Neuroblastoma cell lines LA1-55n, LA1-5s, SH-SY5Y, SHEP, NBL-W-N, NBL-W-S, SK-N-BE2, and NBL-S were obtained and cultured as previously described^24^. SK-N-BE2 and NBL-W-N cells were counted and transferred to T25 cell culture flasks and grown for 24 hours. Cell lines were then treated with either EZH2 inhibitor tazemetostat (ChemieTek CT-EPZ438) at a final concentration of 600nM in Dimethylsulfoxide (DMSO) or DMSO alone for 72 hours. For the cobalt chloride experiments, SK-N-BE2 were cultured in a T75 flask to 50% confluence, and then were treated with H_2_O alone, or cobalt chloride (Sigma-Aldrich C-8661) at concentrations of 10uM, 50uM, or 100uM in H_2_O for 72 hours. 2 x 10^6^ cells were harvested from the flask, pelleted in 1.7ml centrifuge tubes, and resuspended in Puregene Cell Lysis Solution (Qiagen 158113) for DNA extraction. For RNA isolation, cells were harvested from the flask and treated with 2.5ml TRizol Reagent (Invitrogen 15596-026). For the decitabine experiments, SK-N-BE2 wild-type and *TET2* KO cells were seeded at a density of 2.5 × 10⁴ cells per well in 24-well plates and incubated for 24 hours. Cells were then treated with 500nM of Decitabine (Selleckchem S1200) or DMSO alone and monitored in the Cellcyte X™ Live Cell Imaging System. Confluency was measured every 6 hours for a duration of 4 days. Data were processed and analyzed using GraphPad Prism version 10.

### CRISPR-Cas9 knockout of *TET1/TET3* and *TET2/TET3* in SK-N-BE2

CRISPR-Cas9 was utilized to generate *TET1/TET3* and *TET2/TET3* knockout SK-N-BE2 cell lines. Viral transduction of SK-N-BE2 cells was achieved using CRISPR-Cas9 with TET1 VSGH11939-Edit-R Human hEF1a All-in-one lentiviral sgRNA particles (Horizon Discovery), TET2 VSGH11939-Edit-R Human hEF1a All-in-one lentiviral sgRNA particles (Horizon Discovery), and TET3 VSGH12191-Edit-R hEF1a-EGFP All-in-one lentiviral sgRNA particles (Horizon Discovery). A list of sgRNA’s utilized in the current study against *TET1, TET2,* and *TET3* are displayed in **SUPPLEMENTAL TABLE 1, 2, and 3,** respectively. Cells were seeded at a density of 3.0 x 10^5^ per well in a 6-well plate and grown overnight at 37°C in a humidified incubator with 5% CO₂ in RPMI-1640 medium. Double knockout of *TET1* and *TET3* was performed using a single sgRNA targeting *TET1* in combination with three distinct sgRNAs targeting *TET3*. Nine different combinations of sgRNAs were utilized (**SUPPLEMENTAL TABLE 4**). Double knockout of *TET2* and *TET3* was performed using a single sgRNA targeting *TET3* in combination with three distinct sgRNAs targeting *TET2*; a total of six different combinations of sgRNAs were utilized (**SUPPLEMENTAL TABLE 5**). 1µl of 10mg/ml Polybrene (Millipore, TR-1003) was used as a transfection reagent. Finally, the transduced cells were selected using 2µg/ml of puromycin. Cells were further sub-cultured with puromycin selection for seven days. *TET2/TET3* double knockout clones exhibiting reduced 5hmC signal compared to wild-type control by dot blot analysis were selected for further studies (clones #1, 2, and 3).

### DNA Extraction and Dot Blot

Genomic DNA was prepared using Puregene® Cell Core Kit (Qiagen, 1126826) according to the manufacturer’s instructions. DNA was quantified using a DeNovix DS-1 Spectrophtometer and prepared in triplicate with quantities of 2µg, 1µg, and 500ng in a final volume of 600µl. mESC DNA was obtained as a kind gift from the lab of Dr. Ivan Moskowitz and utilized as a positive control. 250ng of mESC DNA was resuspended in nuclease-free H_2_O to a final volume of 300µL. DNA was denatured using 12µl 5M NaOH and incubated at 99°C for 5 minutes. DNA was cooled on ice for 2 min and neutralized with 0.1vol of 6.6M ammonium acetate. Denatured DNA was spotted on a 0.45µm nitrocellulose blotting membrane (Amersham™ Protran™, 10600002) and UV crosslinked at 120mJ/cm2. Blocking of the membrane was done overnight at 4°C using 1% BSA (Sigma, A7030). Probing for global 5hmc was done using 5hmc antibody (Active Motif, 39791) diluted 1:5000 with 1% BSA for 1 hour. Washing of the membrane was performed 3x using Tris-buffered saline with 1% Tween 20 for 10 minutes. Secondary Anti-Rabbit IgG (H+L) antibody (sera care, 5220-0337) was diluted 1:5000 with 1% BSA and the membrane was incubated for 30 minutes. Washing of the membrane was performed 3x using Tris-buffered saline with Tween 20 for 10 minutes. Detection was done using Clarity™ Western ECL Substrate (Bio-Rad, 170-5061) for 5 min and imaging was performed using a ChemiDoc™ Imaging System. The membrane was rinsed with dH_2_O and stained with methylene blue for 5 minutes. The membrane was rinsed with dH_2_O, and colorimetric imaging was performed using ChemiDoc™ Imaging System to visualize DNA control loading.

### Sodium Bisulfite Conversation

SK-N-BE2 knockout and wild type cells were grown to a confluency of 70% in a T25 flask. Cells were harvested and genomic DNA was prepared using Puregene® Cell Core Kit (Qiagen, 1126826) according to the manufacturer’s protocol. DNA was quantified using a DeNovix DS-1 Spectrophtometer. 1µg of DNA in a final volume of 20µl H_2_O was used in a bisulfite conversion reaction using EpiTect® Bisulfite Kit (Qiagen, 59104) according to the manufacture’s protocol. Bisulfite converted DNA was amplified by PCR with primers detailed in **SUPPLEMENTAL TABLE 6**. PCR amplicons were purified using QIAquick PCR Purification Kit (Qiagen, 28106) according to manufacturer’s protocol. Purified PCR products were cloned using CloneJET PCR Cloning Kit (Fisher Scientific, FERK1232) according to the manufacturer’s protocol. 2µl of ligation mixture was added to 50µl of DH5α competent cell prepared in the lab. Cells with ligation mixture were incubated on ice for two minutes and heat shocked at 42°C for 30 seconds. Mixture was incubated on ice again for three minutes followed by the addition of 500µl of LB Broth (Fisher Scientific, BP1426-500) and incubated at 37°C for 1 hour with shaking. Transformed cells were spread on LB Agar plates containing Ampicillin at 100µg/ml. Plates were incubated overnight at 37°C. Bacteria colonies were inoculated with 4ml of LB with 100µg/ml Ampicillin and grown overnight at 37°C with shaking. DNA was prepared from 2ml of inoculated culture using QIAprep Spin Miniprep kit (Qiagen, 27106) according to the manufacturer’s protocol. DNA was quantified using a DeNovix DS-1 Spectrophtometer. Sanger sequencing was performed using sequencing primers pJET 1.2 forward CGACTCACTATAGGGAGAGCGGC (Fisher Scientific, SO501) and reverse pJET 1.2 AAGAACATCGATTTTCCATGGCAG (Fisher Scientific, SO511). Sequencing data was analyzed using Quantification tool for Methylation Analysis (QUMA).

### Chromatin Extraction and ChIP-seq

Chromatin extraction and ChIP-Seq for H3K27me3 was previously performed in the SK-N-BE2 and NBL-W-N cell lines^24^. Using the same procedure previously described, chromatin extraction and ChIP-Seq for H3K27me3 in the NBL-W-S, SHEP, SH-SY5Y, LA1-55n, LA1-5s, and NBL-S cell lines was performed in the current study^24^.

### CUT&RUN Protocol

Cleavage Under Targets & Release Using Nuclease (CUT&RUN) was performed using all reagents contained in the EpiCypher CUTANA ChIC/CUT&RUN Kit Version 3 (EpiCypher #14-1048) according to manufacturer’s instructions. In brief, cells were counted using a Beckman Coulter Vi-CELL XR cell counter to harvest 500,000 cells per CUT&RUN assay. After a brief wash, cells were immobilized to concanavalin A (ConA). Cells were then permeabilized with 0.01% digitonin buffer and bound to 0.5ug of antibody (either IgG control included in EpiCypher kit or anti-H3K27me3, Cell Signaling Technologies #9733) overnight at 4°C. 2µL of the stock SNAP-CUTANA™ K-MetStat panel (EpiCypher #19-1002) was added to each reaction right before adding the primary antibody. After washing off excess antibody, a protein A/G-micrococcal nuclease fusion (pAG-MNase) and calcium chloride (CaCl_2_) were added to all reactions to initiate targeted chromatin digestion. Samples were incubated in this state for 2 hours at 4°C, after which MNase digestion was stopped with an EDTA-containing buffer. Following a 10-minute incubation at 37°C, specifically digested DNA was released into the supernatant of each reaction while whole cells and bulk chromatin remained bound to the ConA beads. As such, the supernatant was isolated from each reaction and DNA was purified using columns included in the kit. 10ng of DNA isolated from CUT&RUN for H3K27me3 was passed directly into the nano-hmC-Seal protocol. Remaining DNA from CUT&RUN for H3K27me3 was subjected to library preparation. Libraries were prepared using the CUTANA CUT&RUN Library Prep Kit (EpiCypher #14-1001) according to manufacturer’s instructions and were sequenced on an Illumina NextSeq 500 in paired-end 150bp mode.

### hmC-Seal Protocol

DNA isolation and hmC-Seal were previously performed in the SK-N-BE2 and NBL-W-N cell lines^24^. Using the same procedure previously described, DNA isolation and hmC-Seal in the NBL-W-S, SHEP, SH-SY5Y, LA1-55n, LA1-5s, and NBL-S cell lines were performed in the current study^24^.

### Nano-hmC-Seal Protocol

DNA from SK-N-BE2 and NBL-W-N cells treated with either tazemetostat or DMSO were isolated using Puregene Cell Kit (Qiagen 158722) according to the manufacturer’s instructions. 100ng of purified genomic DNA was fragmented using KAPA Fragmentation Enzyme and 10x Fragmentation Buffer (Kapa Biosystems KK8514) at 37°C for 30 minutes. An end repair reaction was performed using HyperPrep ERAT Enzyme Mix and ERAT Buffer (Kapa Biosystems KK8514) for 30 minutes at 65°C. DNA adapter ligation was done using NEXTflex® DNA Barcodes (Perkin Elmer NOVA 514104), KAPA Hyper Plus DNA Ligase and Buffer (Kapa Biosystems KK8514) at 20°C for 1 hour.

To determine H3K27me3-associated 5-hmC, DNA from CUT&RUN for H3K27me3 conducted on untreated SK-N-BE2 and NBL-W-N cells underwent an end repair using HyperPrep ERAT Enzyme Mix and ERAT Buffer (Kapa Biosystems KK8514) for 30 minutes at 20°C then 30 minutes at 65°C. DNA adapter ligation was done using NEXTflex® DNA Barcodes (Perkin Elmer NOVA 514104), KAPA Hyper Plus DNA Ligase and Buffer (Kapa Biosystems KK8514) at 20°C for 4 hours.

All adapter ligation reactions were purified using DNA Clean & Concentrator™-5 Kit (Zymo D4014) according to the manufacturer’s instructions and eluted in 20µl of nuclease free H_2_O. Glucosylation reactions were performed in a 26µl solution containing purified DNA with UDP-Azide-Glucose (Active Motif 55020), T4 beta-glucosyltransferase and 10X EPi Buffer (Fisher Scientific FEREO0831) and incubated at 37°C for 2 hours. Separate control glucosylation reactions (sham-nano-Seal) were done for CUT&RUN SK-N-BE2 and NBL-W-N purified DNA that did not contain T4 beta-glucosyltransferase. The DNA was purified using DNA Clean & Concentrator™-5 Kit (Zymo D4014) according to the manufacturer’s instructions and eluted in 30µl of nuclease free water. The modified DNA was added to 1µl 4.5mM DBCO-PEG4 Biotin (Sigma 760749) and incubated at 37°C for 2 hours. The DNA was purified using DNA Clean & Concentrator™-5 Kit (Zymo D4014) according to the manufacturer’s instructions and eluted in 30µl of nuclease free water. Invitrogen™ Dynabeads™ MyOne™ Streptavidin C1 beads (Fisher Scientific 65-001) were washed with Buffer 1 containing 5mM Tris-HCl pH 7.5, 0.5mM EDTA, 1M NaCl, and 2% Tween 20. A magnetic rack was used to remove the solution from the beads. The beads were then incubated for 30 minutes at room temperature with Invitrogen™ UltraPure™ Salmon Sperm DNA Solution (Fisher Scientific 15-632-011) and Buffer 1 with top-over-end-rotation. Beads were washed again with Buffer 1 and resuspended in Binding Buffer containing 10mM Tris-HCl pH 7.5, 1mM EDTA, 2M NaCl, and 4% Tween. Beads were added to purified DNA and incubated at room temperature for 30 minutes with top-over-end rotation. Two washes with Buffer 1 and two washes of each of the following buffers were performed on the beads. Buffer 2 contained 5mM Tris pH 7.5, 0.5mM EDTA, and 2% Tween 20. Buffer 3 contained 5mM Tris-HCl pH 9.0, 0.5mM EDTA, 1M NaCl, and 2% Tween 20. Buffer 4 contained 5mM Tris-HCl pH 9.0, 0.5mM EDTA, and 2% Tween 20. Following the final wash, beads were resuspended in 23.8µl of nuclease free H_2_O. Captured DNA fragments were amplified with 15 cycles of PCR with KAPA HiFi Hot Start Ready Mix (2x) (Kapa Biosystems KK8514). The PCR products were purified using AMPure XP beads (Beckman Coulter A63880). DNA libraries were sequenced on an Illumina NovaSeq 6000 in paired end 50bp mode.

### Western Blot Protocol

For Western blotting, SK-N-BE2 and NBL-W-N cell pellets were collected. Cell pellets were washed with phosphate buffered saline (PBS, pH 7.4). Pellets were then suspended and boiled for 5 minutes in lysis buffer containing 50mM Tris-HCl pH 6.8 and 2% SDS. BCA Protein Assay Reagent (Pierce) was utilized to determine protein concentration. Total protein concentrations across samples were normalized. Protein lysates were mixed with Laemmli Sample Buffer and beta-mercaptoethanol and then boiled for 5 minutes. 30µg of protein from each sample were loaded onto a Criterion TGX Stain-Free Gel (Bio-Rad Catalog #5678124) and electrophoresed and transferred to a nitrocellulose membrane (Amersham). Blocking, staining with primary and secondary antibodies, and blot development were performed as previously described ^60^. Primary antibodies used in the current study include 1:1,000 anti-H3K27me3 (Cell Signaling Technologies #9733), 1:1,000 anti-H3 (Active Motif #39763), 1:1000 anti-TET1 (Santa Cruz #sc-293186), 1:1000 anti-TET2 (Cell Signaling #18950), and 1:1000 anti-TET3 (ProSci #7013). Secondary antibodies used in the current study include 1:10,000 anti-rabbit (KPL #5220-0337) and 1:10,000 anti-mouse (KPL #5220-0341).

### RNA sequencing and RNA library construction

RNA isolation, library preparation, and sequencing was performed as previously described^24^.

### Sequencing data processing, analysis, and visualization

fastqc (version 0.11.5) with default settings was used to assess read quality for all included sequencing data^49^. For cell line ChIP-Seq, CUT&RUN, and 5-hmC sequencing data, raw reads with a minimum Phred score of 33 were filtered with Trimmomatic (version 0.36). Using Bowtie2 (version 2.3.0) with default settings, filtered reads were aligned to GRCh38^50,51^. Picard (version 2.18.29) with default settings was utilized to eliminate duplicate reads^52^. Raw 5-hmC counts were loaded into DESeq2 for differential expression analysis where a top 10% Log2FC and FDR < 0.05 was considered significant. In ChIP-Seq experiments, H3K27me3 peaks were defined by MACS2 (version 2.1.0) using the --broad flag^61^. deepTools (version 2.0) “bamCompare” was utilized to define relative ChIP occupancy signal for H3K27me3 relative to input control samples^62^. H3K27me3 enriched peaks were defined as peaks with greater than 2 log_2_fold increase in ChIP signal compared to input^62^. Peaks were then assigned to genes if they fell within 3kb of the TSS. For hmC-Seal, CUT&RUN, and CUT&RUN followed by nano-hmC-Seal, SICER2 with default settings was utilized to identify native 5-hmC, H3K27me3, and H3K27me3-associated-5hmC peaks, respectively^63^.

H3K27me3 and H3K27me3-associated-5hmC peaks were defined as those having at least 2-fold increase in signal compared to input. Native 5-hmC, H3K27me3, and H3K27me3-associated-5hmC peaks were then assigned to genes if they fell within 3kb of the TSS. deepTools (version 2.0) was utilized to generate bigwig files normalized to reads per kilobase million (RPKM) and visualize ChIP-Seq, CUT&RUN, and 5-hmC profiles^62^. For CUT&RUN samples with spike-in included, the scaling factor was determined by dividing the total percentage of human reads by the total percentage of barcode reads, and then dividing again by 100. Scaling factors were applied to bigwig files through the “--scaleFactor” flag in deepTools. Overlap between gene sets was assessed with the hypergeometric t-test.

For RNA-Seq data, raw reads with a minimum phred score of 33 were filtered with Trimmomatic (version 0.36). STAR-aligner (version 2.6.1d) was then used to align reads to GRCh38^50,64^. featureCounts of subread (version 1.5.3) was used to count aligned reads across exons^53^. Raw counts were loaded into DESeq2 for differential expression analysis where a 10% Log2FC and FDR < 0.05 was considered significant, and were also separately converted to transcripts per million (TPM)^55^.

### Survival analysis in neuroblastoma tumors

Phenotypic and expression data for 498 neuroblastoma tumors (SEQC-NB cohort) and a separate cohort of 105 neuroblastoma tumors (GSE73517) were downloaded from R2^35,36,65^. Gene set variance analysis (GSVA) was performed using the *GSVA* package (version 1.46.0) on included tumor samples using previously published *MYCN* signatures and 5-hmC/H3K27me3 co-enriched genes, as identified in the current study^37,38^. To assess the power of our 5-hmC/H3K27me3 signature scores to predict overall survival, area under the receiver operator curve (AUROC) analyses were performed using the pROC package (version 1.18.0)^66^. To maximize both sensitivity and specificity, optimal 5-hmC/H3K27me3 co-enriched gene signature GSVA score cutoff (−0.15) was determined in the HR-only subset of the SEQC-NB cohort (n=175 tumors) using the “thresholds = “best”” parameter of the pROC package. A GSVA score cutoff of -0.15 was utilized to tumors with “5-hmC/H3K27me3 low expression” or “5-hmC/H3K27me3 high expression” across both included datasets. EFS and OS were determined using the Kaplan-Meier method. Log-rank test was utilized to assess differences in survival between groups. “survival” package (version 3.5-3) was used to perform survival analyses^67^.

## ACKNOWLEDGEMENTS

This work was supported in part by the Burroughs Wellcome Fund Early Scientific Training Program to Prepare for Research Excellence Post-Graduation (BEST-PREP; MC), University of Chicago Pritzker School of Medicine Summer Research Program (SRP; MC), the Alex’s Lemonade Stand Foundation (MAA), and a kind gift from Barry & Kimberly Fields (CH). Also, supported by the National Institutes of Health R37CA262781 (MAA), R00CA234424 (AP) and the Ludwig Center at the University of Chicago (CH). CH is a Howard Hughes Medical Institute Investigator. Higher performance computing used is funded by the Biological Sciences Division at the University of Chicago with additional funding provided by the Institute for Translational Medicine, CTSA grant number UL1TR000430. The contents are solely the responsibility of the authors and do not necessarily represent the official views of the NIH.

## AUTHOR CONTRIBUTIONS

AP, MAA, and MC designed the study and wrote the manuscript. VG, KM, GLL, YX, JM, RB, MC, and MP performed experiments. MC and SV retrieved sequencing data and performed bioinformatics and statistical analyses. MC, VG, AC, CH, AP, MAA assisted with data interpretation. MAA provided clinical insights on the project. All authors reviewed, edited, and approved the submitted manuscript.

## DATA AVAILABILITY

ChIP-Seq, CUT&RUN, 5-hmC, and RNA-Seq raw and processed data included in the current study are available at GSE294304, GSE294307, GSE294308.

## CODE AVAILABILITY

The underlying code for this study is not publicly available but may be made available on reasonable request from the corresponding author.

## REFERENCES

1. Pinto, N. R. et al. Advances in Risk Classification and Treatment Strategies for Neuroblastoma. J. Clin. Oncol. 33, 3008–3017 (2015).

2. Althoff, K. et al. A Cre-conditional MYCN-driven neuroblastoma mouse model as an improved tool for preclinical studies. Oncogene 34, 3357–3368 (2015).

3. Qiu, B. & Matthay, K. K. Advancing therapy for neuroblastoma. Nat. Rev. Clin. Oncol. 19, 515–533 (2022).

4. Park, J. R. et al. Effect of Tandem Autologous Stem Cell Transplant vs Single Transplant on Event-Free Survival in Patients With High-Risk Neuroblastoma: A Randomized Clinical Trial. JAMA 322, 746–755 (2019).

5. Zimmerman, M. W. et al. MYC Drives a Subset of High-Risk Pediatric Neuroblastomas and Is Activated through Mechanisms Including Enhancer Hijacking and Focal Enhancer Amplification. Cancer Discov. 8, 320–335 (2018).

6. Fetahu, I. S. & Taschner-Mandl, S. Neuroblastoma and the epigenome. Cancer Metastasis Rev. 40, 173–189 (2021).

7. Pugh, T. J. et al. The genetic landscape of high-risk neuroblastoma. Nat. Genet.45, 279–284 (2013).

8. Durinck, K. & Speleman, F. Epigenetic regulation of neuroblastoma development. Cell Tissue Res. 372, 309–324 (2018).

9. Tsubota, S. et al. PRC2-Mediated Transcriptomic Alterations at the Embryonic Stage Govern Tumorigenesis and Clinical Outcome in MYCN-Driven Neuroblastoma. Cancer Res. 77, 5259–5271 (2017).

10. Chen, L. et al. CRISPR-Cas9 screen reveals a MYCN-amplified neuroblastoma dependency on EZH2. J. Clin. Invest. 128, 446–462 (2018).

11. Piunti, A. & Shilatifard, A. Epigenetic balance of gene expression by Polycomb and COMPASS families. Science 352, aad9780 (2016).

12. Piunti, A. & Shilatifard, A. The roles of Polycomb repressive complexes in mammalian development and cancer. Nat. Rev. Mol. Cell Biol. 22, 326–345 (2021).

13. Margueron, R. & Reinberg, D. The Polycomb complex PRC2 and its mark in life. Nature 469, 343–349 (2011).

14. Zhang, H. et al. TET1 is a DNA-binding protein that modulates DNA methylation and gene transcription via hydroxylation of 5-methylcytosine. Cell Res. 20, 1390–1393 (2010).

15. Applebaum, M. A. et al. 5-Hydroxymethylcytosine Profiles Are Prognostic of Outcome in Neuroblastoma and Reveal Transcriptional Networks That Correlate With Tumor Phenotype. *JCO Precis*. Oncol. 3, (2019).

16. Rasmussen, K. D. & Helin, K. Role of TET enzymes in DNA methylation, development, and cancer. Genes Dev. 30, 733–750 (2016).

17. Applebaum, M. A. et al. 5-Hydroxymethylcytosine Profiles in Circulating Cell-Free DNA Associate with Disease Burden in Children with Neuroblastoma. Clin. Cancer Res. Off. J. Am. Assoc. Cancer Res. 26, 1309–1317 (2020).

18. Neri, F. et al. Genome-wide analysis identifies a functional association of Tet1 and Polycomb repressive complex 2 in mouse embryonic stem cells. Genome Biol. 14, R91 (2013).

19. Wu, H. et al. Genome-wide analysis of 5-hydroxymethylcytosine distribution reveals its dual function in transcriptional regulation in mouse embryonic stem cells. Genes Dev. 25, 679–684 (2011).

20. Wu, H. et al. Dual functions of Tet1 in transcriptional regulation in mouse embryonic stem cells. Nature 473, 389–393 (2011).

21. Pastor, W. A. et al. Genome-wide mapping of 5-hydroxymethylcytosine in embryonic stem cells. Nature 473, 394–397 (2011).

22. Otte, J., Dyberg, C., Pepich, A. & Johnsen, J. I. MYCN Function in Neuroblastoma Development. Front. Oncol. 10, 624079 (2020).

23. Lachmann, A. et al. ChEA: transcription factor regulation inferred from integrating genome-wide ChIP-X experiments. Bioinforma. Oxf. Engl. 26, 2438–2444 (2010).

24. Chennakesavalu, M. et al. 5-Hydroxymethylcytosine Profiling of Cell-Free DNA Identifies Bivalent Genes That Are Prognostic of Survival in High-Risk Neuroblastoma. *JCO Precis*. Oncol. 8, e2300297 (2024).

25. Williams, K. et al. TET1 and hydroxymethylcytosine in transcription and DNA methylation fidelity. Nature 473, 343–348 (2011).

26. Han, D. et al. A Highly Sensitive and Robust Method for Genome-wide 5hmC Profiling of Rare Cell Populations. Mol. Cell 63, 711–719 (2016).

27. Wang, C. et al. EZH2 Mediates epigenetic silencing of neuroblastoma suppressor genes CASZ1, CLU, RUNX3, and NGFR. Cancer Res. 72, 315–324 (2012).

28. Brinkman, A. B. et al. Sequential ChIP-bisulfite sequencing enables direct genome-scale investigation of chromatin and DNA methylation cross-talk. Genome Res. 22, 1128–1138 (2012).

29. Mabe, N. W. et al. Transition to a mesenchymal state in neuroblastoma confers resistance to anti-GD2 antibody via reduced expression of ST8SIA1. *Nat*. Cancer 3, 976–993 (2022).

30. Sengupta, S. et al. Mesenchymal and adrenergic cell lineage states in neuroblastoma possess distinct immunogenic phenotypes. *Nat*. Cancer 3, 1228–1246 (2022).

31. Tahiliani, M. et al. Conversion of 5-methylcytosine to 5-hydroxymethylcytosine in mammalian DNA by MLL partner TET1. Science 324, 930–935 (2009).

32. Muñoz-Sánchez, J. & Chánez-Cárdenas, M. E. The use of cobalt chloride as a chemical hypoxia model. J. Appl. Toxicol. JAT 39, 556–570 (2019).

33. Ito, S. et al. Role of Tet proteins in 5mC to 5hmC conversion, ES-cell self-renewal and inner cell mass specification. Nature 466, 1129–1133 (2010).

34. Pastor, W. A., Aravind, L. & Rao, A. TETonic shift: biological roles of TET proteins in DNA demethylation and transcription. Nat. Rev. Mol. Cell Biol. 14, 341–356 (2013).

35. Henrich, K.-O. et al. Integrative Genome-Scale Analysis Identifies Epigenetic Mechanisms of Transcriptional Deregulation in Unfavorable Neuroblastomas. Cancer Res. 76, 5523–5537 (2016).

36. Wang, C. et al. The concordance between RNA-seq and microarray data depends on chemical treatment and transcript abundance. Nat. Biotechnol. 32, 926–932 (2014).

37. Hänzelmann, S., Castelo, R. & Guinney, J. GSVA: gene set variation analysis for microarray and RNA-seq data. BMC Bioinformatics 14, 7 (2013).

38. Valentijn, L. J. et al. Functional MYCN signature predicts outcome of neuroblastoma irrespective of MYCN amplification. Proc. Natl. Acad. Sci. U. S. A. 109, 19190–19195 (2012).

39. Li, W. et al. 5-Hydroxymethylcytosine signatures in circulating cell-free DNA as diagnostic biomarkers for human cancers. Cell Res. 27, 1243–1257 (2017).

40. Reddington, J. P. et al. Redistribution of H3K27me3 upon DNA hypomethylation results in de-repression of Polycomb target genes. Genome Biol. 14, R25 (2013).

41. Jermann, P., Hoerner, L., Burger, L. & Schübeler, D. Short sequences can efficiently recruit histone H3 lysine 27 trimethylation in the absence of enhancer activity and DNA methylation. Proc. Natl. Acad. Sci. U. S. A. 111, E3415–3421 (2014).

42. Weigert, R. et al. Dynamic antagonism between key repressive pathways maintains the placental epigenome. Nat. Cell Biol. 25, 579–591 (2023).

43. Richard Albert, J., et al. DNA methylation shapes the Polycomb landscape during the exit from naive pluripotency. Nat. Struct. Mol. Biol. 32, 346–357 (2025).

44. Qadeer, Z. A. et al. ATRX In-Frame Fusion Neuroblastoma Is Sensitive to EZH2 Inhibition via Modulation of Neuronal Gene Signatures. Cancer Cell 36, 512–527.e9 (2019).

45. Yamagishi, M. et al. Mechanisms of action and resistance in histone methylation-targeted therapy. Nature 627, 221–228 (2024).

46. Mohn, F. et al. Lineage-specific polycomb targets and de novo DNA methylation define restriction and potential of neuronal progenitors. Mol. Cell 30, 755–766 (2008).

47. Jansky, S. et al. Single-cell transcriptomic analyses provide insights into the developmental origins of neuroblastoma. Nat. Genet. 53, 683–693 (2021).

48. Kameneva, P. et al. Single-cell transcriptomics of human embryos identifies multiple sympathoblast lineages with potential implications for neuroblastoma origin. Nat. Genet. 53, 694–706 (2021).

49. Andrews, S. FastQC: a quality control tool for high throughput sequence data. http://www.bioinformatics.babraham.ac.uk/projects/fastqc/.

50. Bolger, A. M., Lohse, M. & Usadel, B. Trimmomatic: a flexible trimmer for Illumina sequence data. Bioinforma. Oxf. Engl. 30, 2114–2120 (2014).

51. Langmead, B. & Salzberg, S. L. Fast gapped-read alignment with Bowtie 2. Nat. Methods 9, 357–359 (2012).

52. Picard: A set of command line tools (in Java) for manipulating high-throughput sequencing (HTS) data and formats such as SAM/BAM/CRAM and VCF. http://broadinstitute.github.io/picard/.

53. Liao, Y., Smyth, G. K. & Shi, W. The Subread aligner: fast, accurate and scalable read mapping by seed-and-vote. Nucleic Acids Res. 41, e108–e108 (2013).

54. Tang, Y., Horikoshi, M. & Li, W. ggfortify: Unified Interface to Visualize Statistical Result of Popular R Packages. R J. 8, (2016).

55. Love, M. I., Huber, W. & Anders, S. Moderated estimation of fold change and dispersion for RNA-seq data with DESeq2. Genome Biol. 15, 550 (2014).

56. EnhancedVolcano: Publication-ready volcano plots with enhanced colouring and labeling.

57. Gu, Z., Eils, R. & Schlesner, M. Complex heatmaps reveal patterns and correlations in multidimensional genomic data. Bioinforma. Oxf. Engl. 32, 2847–2849 (2016).

58. Chen, E. Y. et al. Enrichr: interactive and collaborative HTML5 gene list enrichment analysis tool. BMC Bioinformatics 14, 128 (2013).

59. Raudvere, U. et al. g:Profiler: a web server for functional enrichment analysis and conversions of gene lists (2019 update). Nucleic Acids Res. 47, W191–W198 (2019).

60. Yu, G., Wang, L.-G., Han, Y. & He, Q.-Y. clusterProfiler: an R package for comparing biological themes among gene clusters. Omics J. Integr. Biol. 16, 284–287 (2012).

61. Zhang, Y. et al. Model-based analysis of ChIP-Seq (MACS). Genome Biol. 9, R137 (2008).

62. Ramírez, F., Dündar, F., Diehl, S., Grüning, B. A. & Manke, T. deepTools: a flexible platform for exploring deep-sequencing data. Nucleic Acids Res. 42, W187–191 (2014).

63. Xu, S., Grullon, S., Ge, K. & Peng, W. Spatial clustering for identification of ChIP-enriched regions (SICER) to map regions of histone methylation patterns in embryonic stem cells. Methods Mol. Biol. Clifton NJ 1150, 97–111 (2014).

64. Dobin, A. et al. STAR: ultrafast universal RNA-seq aligner. Bioinforma. Oxf. Engl. 29, 15–21 (2013).

65. ‘R2: Genomics Analysis and Visualization Platform (http://r2.amc.nl)’.

66. Robin, X. et al. pROC: an open-source package for R and S+ to analyze and compare ROC curves. BMC Bioinformatics 12, 77 (2011).

67. Therneau, Terry. A Package for Survival Analysis in R. (2023).

