## Supplemental Figures and Tables for "5-hydroxymethylcytosine deposition mediates Polycomb Repressive Complex 2 function in *MYCN*-amplified neuroblastoma"

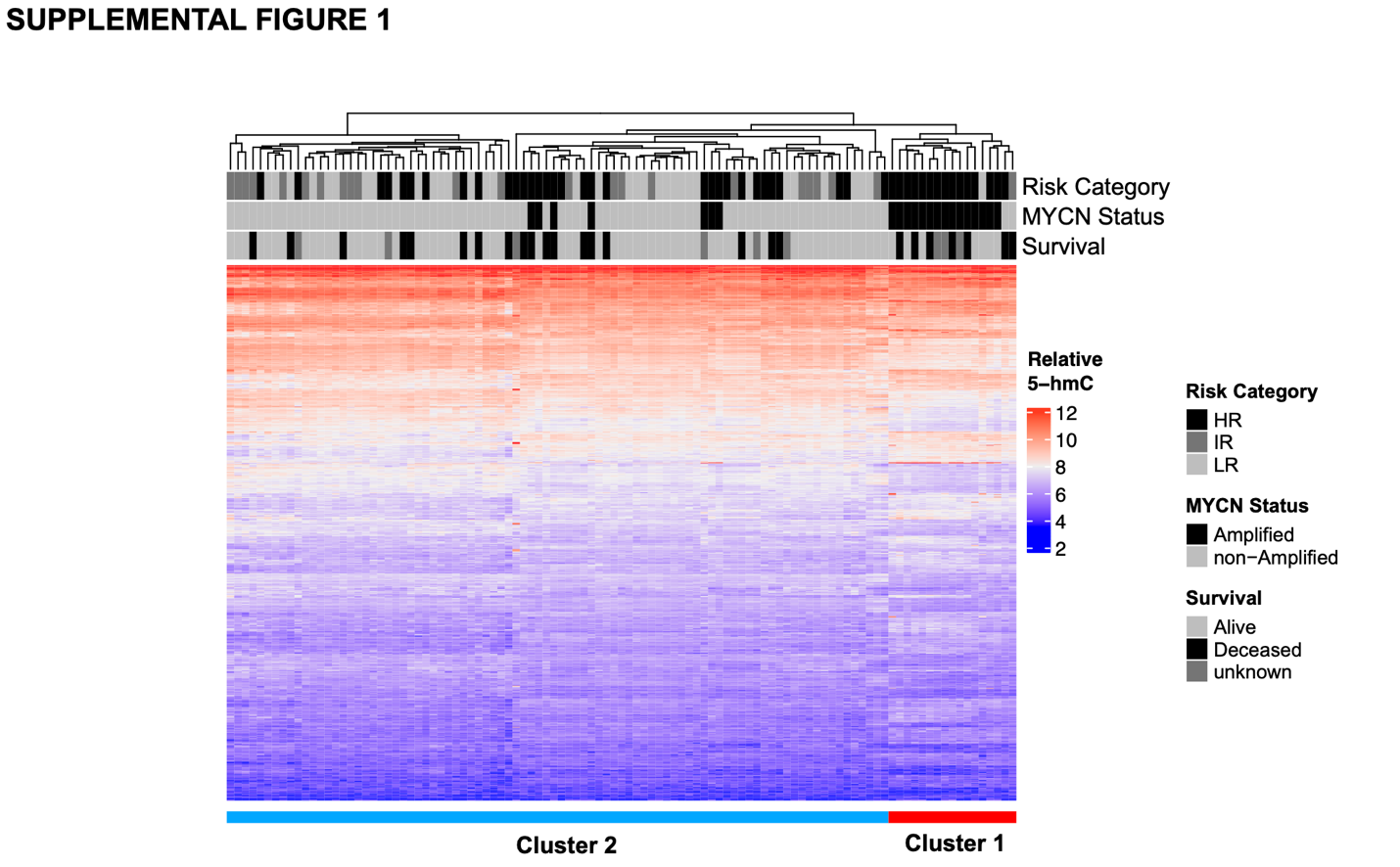

**SUPPLEMENTAL FIGURE 1: Unsupervised clustering of neuroblastoma patient tumor derived 5-hmC profiles**. Unsupervised hierarchical clustering of neuroblastoma patient tumor derived 5-hmC profiles by the top 10% of significantly differentially hydroxymethylated genes (n=901) identified in a comparison of *MYCN-*amplified (n=22) and non-amplified tumors (n=83). Two distinct clusters of patients, correlating to *MYCN*-amplification status, are identified, as noted at the bottom of the figure.

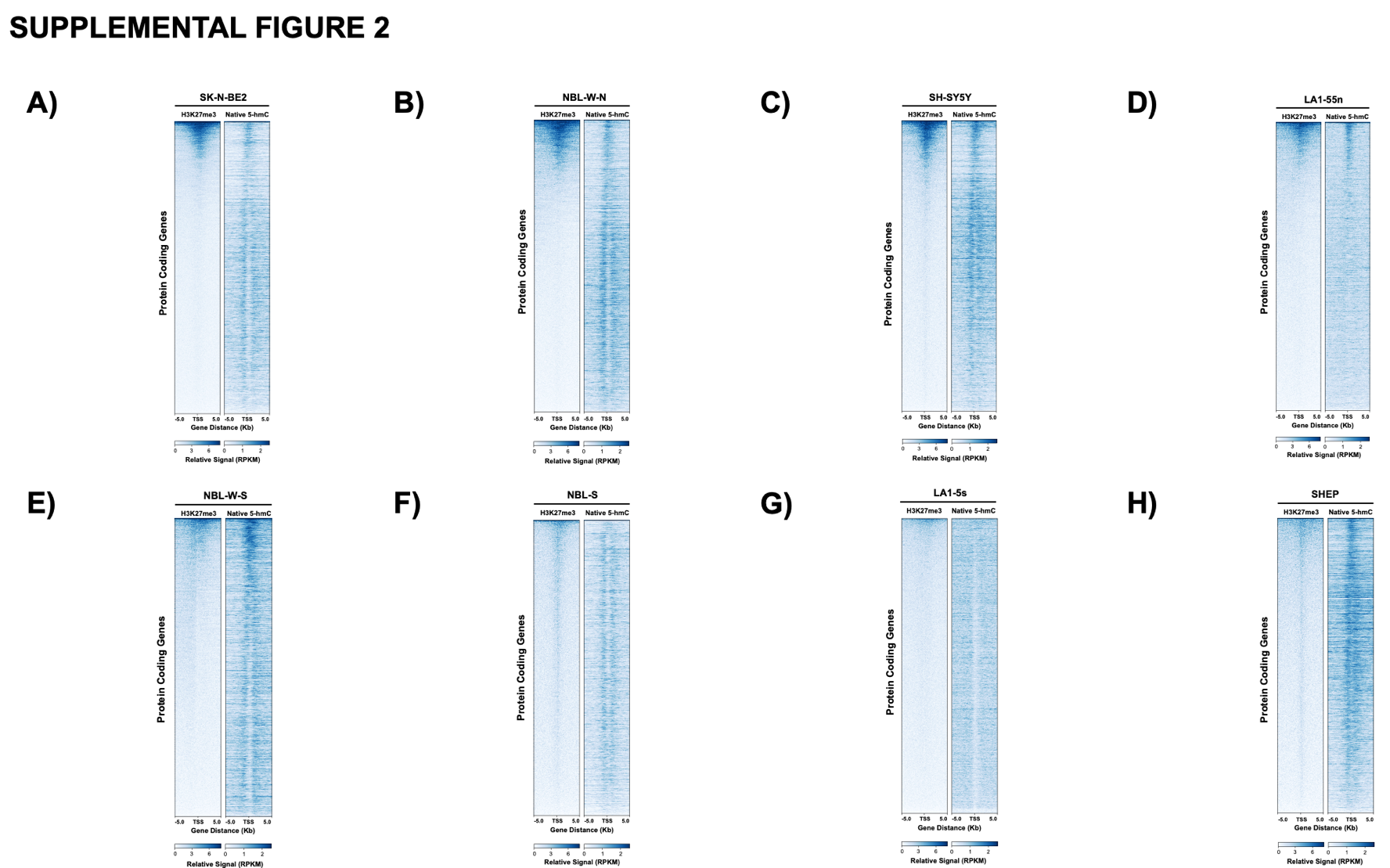

**SUPPLEMENTAL FIGURE 2: Genome-wide H3K27me3 and native 5-hmC profiles across a panel of eight neuroblastoma cell lines.** H3K27me3 and native 5-hmC profiles of protein coding genes, in the **A)** SK-N-BE2, **B)** NBL-W-N, **C)** SH-SY5Y, **D)** LA1-55n, **E)** NBL-W-S, **F)** NBL-S, **G)** LA1-5s, and **H)** SHEP cell lines. Profiles are ordered by relative H3K27me3 signal within each cell line.

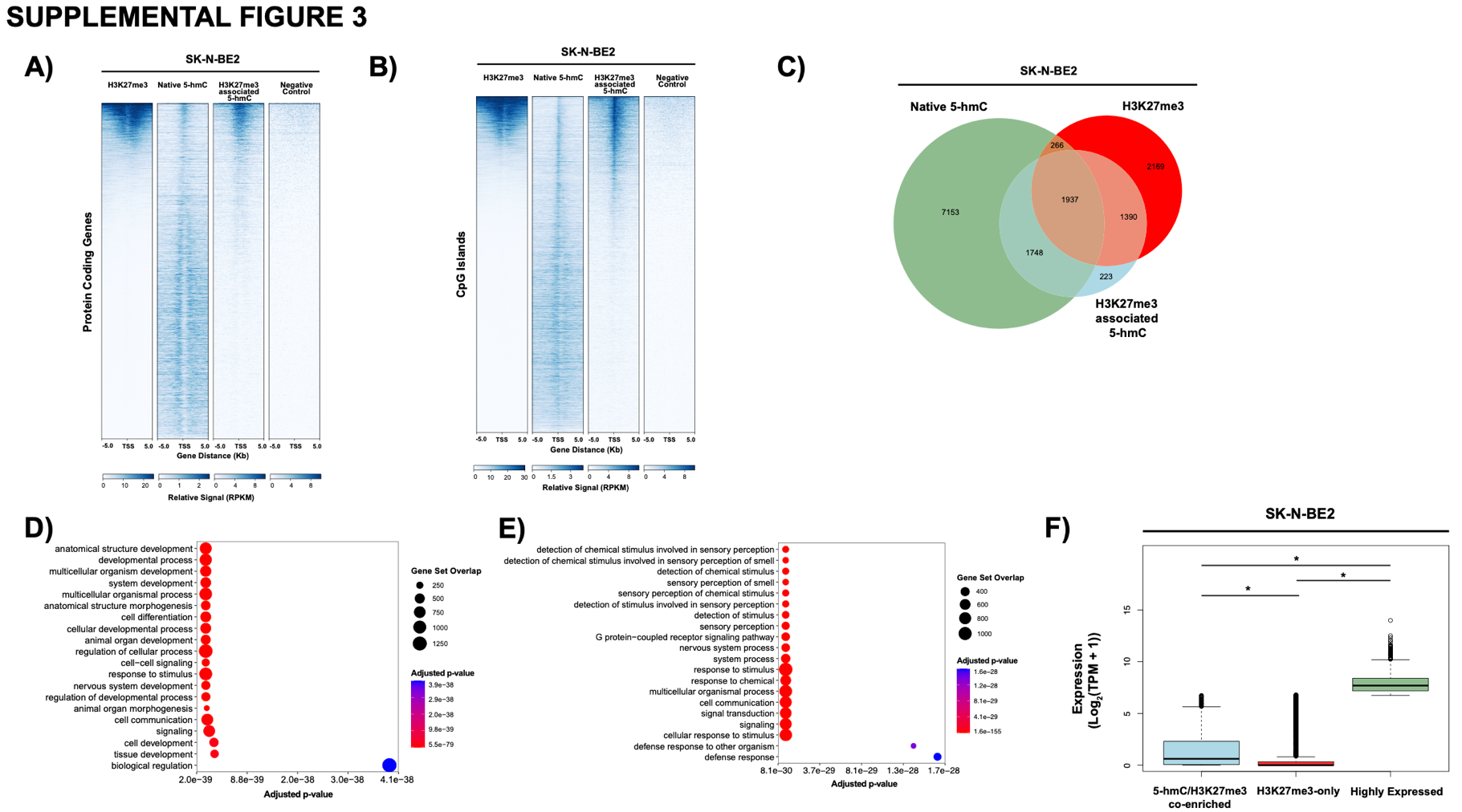

**SUPPLEMENTAL FIGURE 3: Transcription start site 5-hmC and H3K27me3 directly co-localize at the nucleosomal level in SK-N-BE2. A)** H3K27me3, native 5-hmC, H3K27me3 associated 5-hmC, and negative control profiles of protein coding genes in the SK-N-BE2 cell line. **B)** H3K27me3, native 5-hmC, H3K27me3 associated 5-hmC, and negative control profiles of CpG islands in the SK-N-BE2 cell line. **C)** Euler diagram showing overlap of genes with significant enrichment of native 5-hmC, H3K27me3, and H3K27me3 associated 5-hmC. **D)** Expression across 5-hmC/H3K27me3 enriched, H3K27me3-only enriched, and highly expressed (top 2000 genes by TPM) in SK-N-BE2. * denotes p-value < 0.05. P-value determined through two-tailed t-test. **E)** Top 20 significant (FDR<0.05) Gene Ontology: Biological Processes (GO:BP) enriched in genes (n=1,937) with 5-hmC/H3K27me3 co-enrichment in SK-N-BE2. **F)** Top 20 significant (FDR<0.05) GO:BP enriched in genes (n=2,169) with H3K27me3-only enrichment in SK-N-BE2.

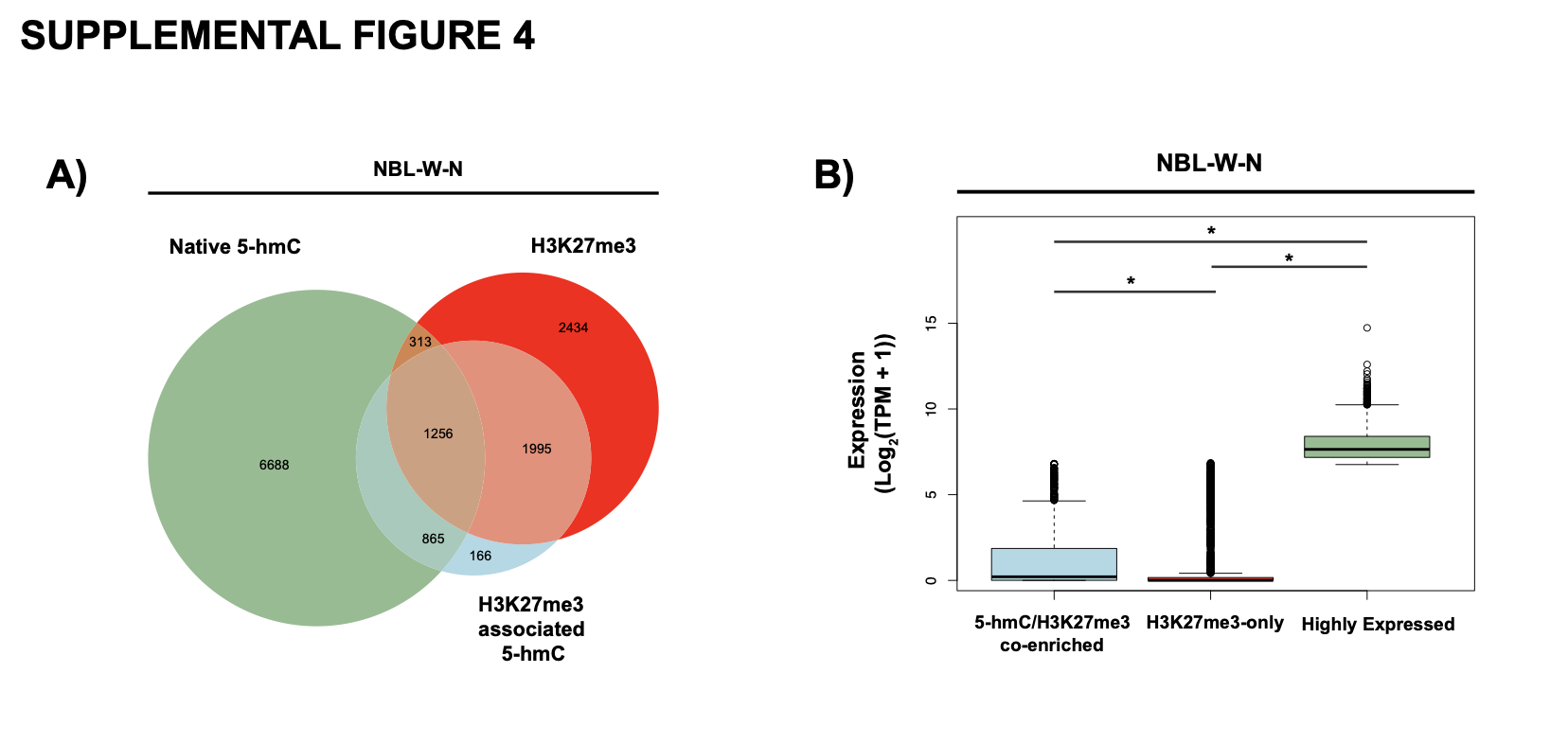

**SUPPLEMENTAL FIGURE 4: Characterization of 5-hmC/H3K27me3 co-enriched and H3K27me3-only enriched genes in the NBL-W-N cell line.** **A)** Euler diagram showing overlap of genes with significant enrichment of native 5-hmC, H3K27me3, and H3K27me3 associated 5-hmC. **B)** Expression across 5-hmC/H3K27me3 enriched, H3K27me3-only enriched, and highly expressed (top 2000 genes by TPM) in NBL-W-N. * denotes p-value < 0.05. P-value determined through two-tailed t-test.

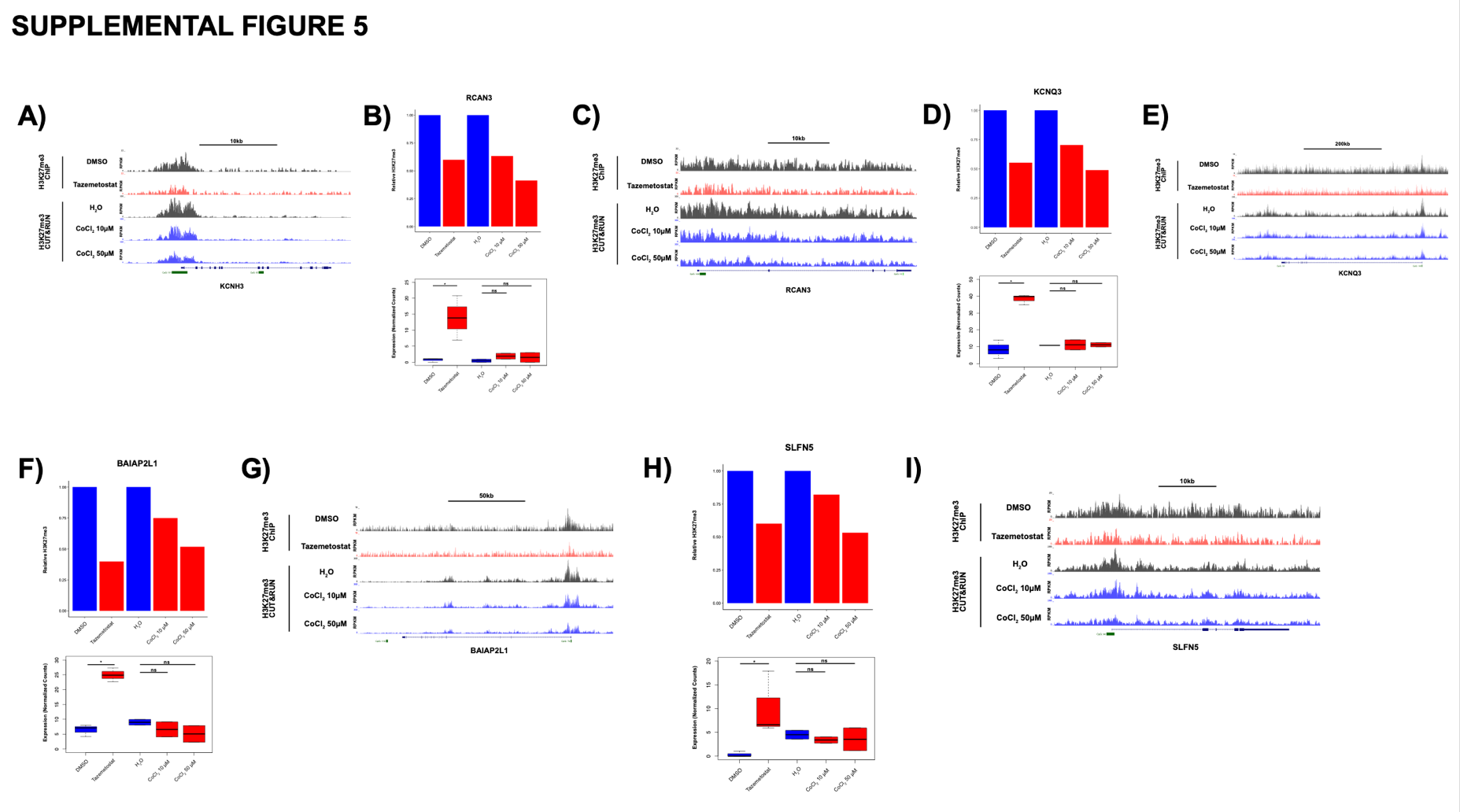

**SUPPLEMENTAL FIGURE 5: Chemical inhibition of 5-hmC deposition with cobalt chloride results in a loss of H3K27me3 but does not result in transcriptional activation of 5-hmC/H3K27me3 co-enriched genes.** H3K27me3 tracks (normalized to RPKM) are shown for **A)** *KCNH3*, **C)** *RCAN3,* **E)** *KCNQ3*, **G)** *BAIAP2L1*, and **I)** *SLFN5* in DMSO, tazemetostat, H_2_O, and CoCl_2_ treated SK-N-BE2 cells. Relative H3K27me3 signal of **B)** *RCAN3,* **D)** *KCNQ3,* **F)** *BAIAP2L1*, **H)** *SLFN5* is shown for DMSO, tazemetostat, H_2_O, and CoCl_2_ treated SK-N-BE2 cells. Tazemetostat treated cells were normalized to DMSO treated cells, and CoCl_2_ treated cells were normalized to H_2_O treated cells. Expression, in DeSeq2 normalized counts, of **B)** *RCAN3,* **D)** *KCNQ3,* **F)** *BAIAP2L1*, **H)** *SLFN5* is shown for DMSO, tazemetostat, H_2_O, and CoCl_2_ treated SK-N-BE2 cells.

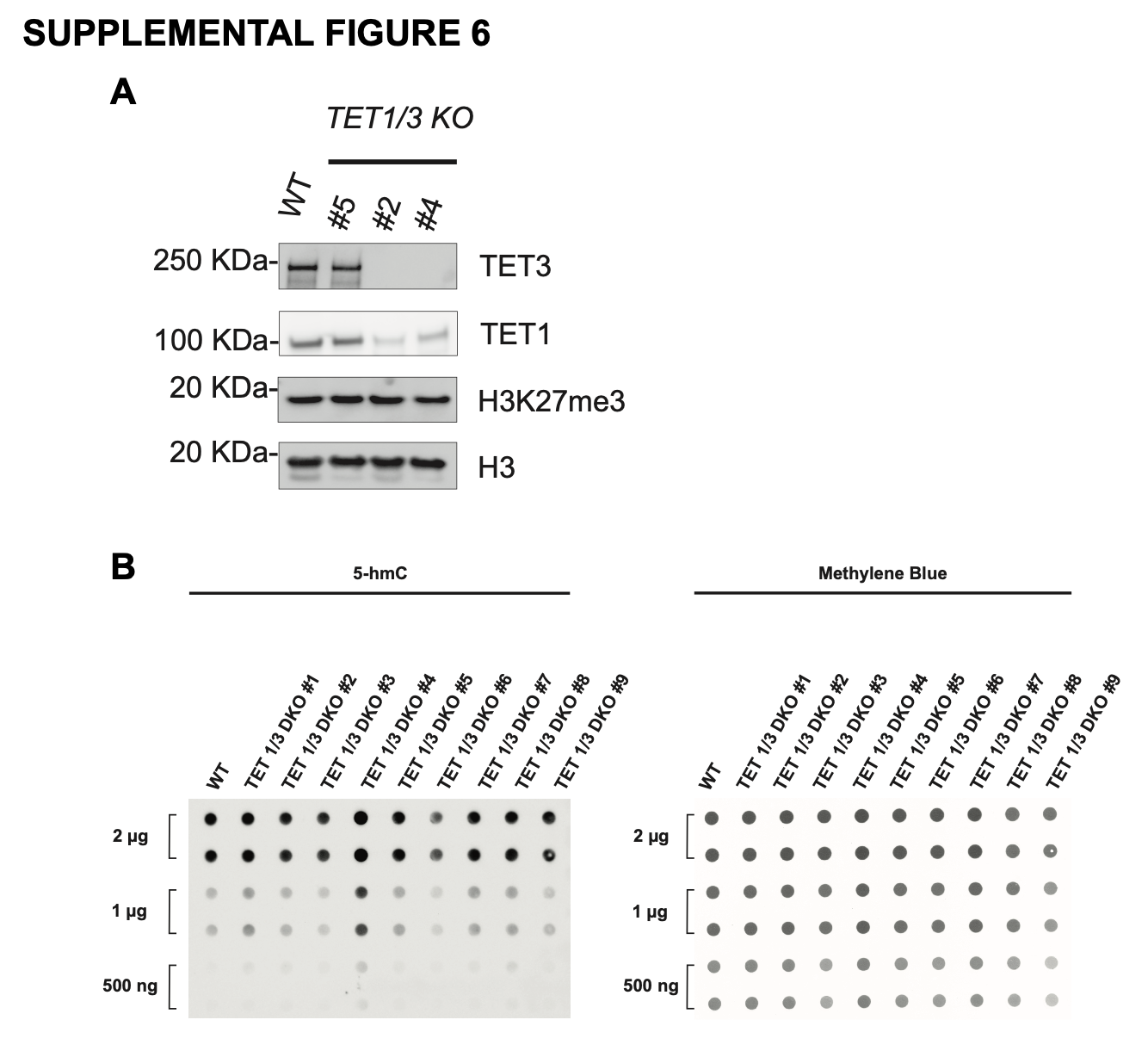

**SUPPLEMENTAL FIGURE 6: TET1/3 double knockout does not lead to global changes in the level of 5-hmC in SK-N-BE2**. **A)** Western blot showing relative levels of TET3, TET1, H3K27me3, and total H3 in SK-N-BE2 wild type and *TET1/3* CRISPR-Cas9 mediated double knockout (DKO) cells. **B)** Dot-blot showing relative 5-hmC (left) and total DNA by methylene blue stain (right) in WT and *TET1/3* DKO SK-N-BE2 cells. Dot blot was run with three different quantities of DNA as indicated.

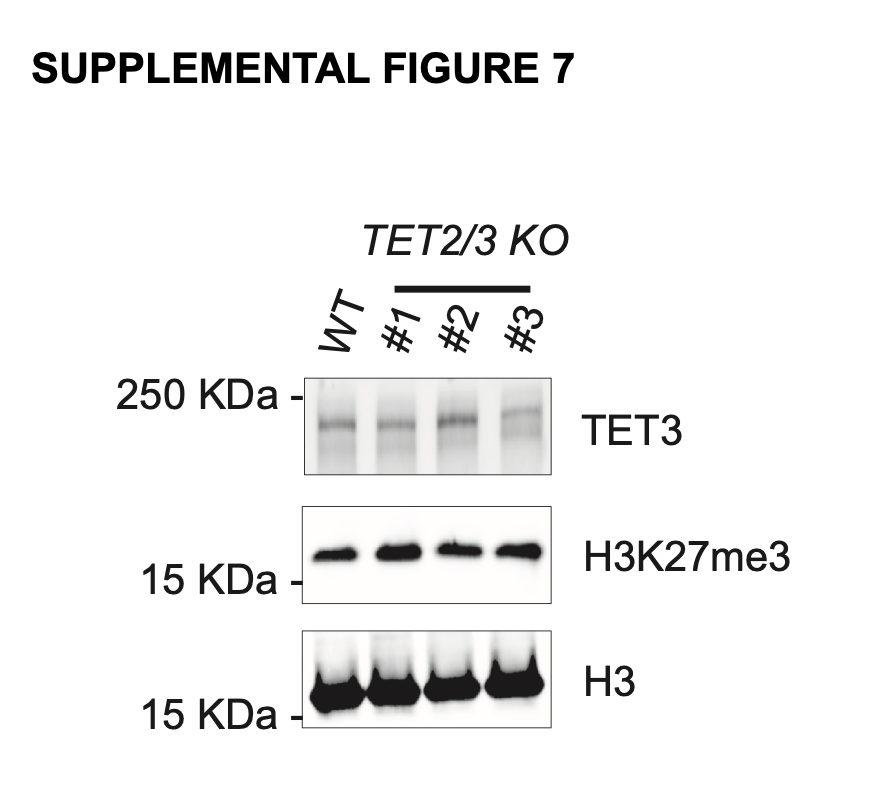

**SUPPLEMENTAL FIGURE 7: *TET2/3* double knockout does not lead to loss of TET3 protein expression in SK-N-BE2**. **A)** Western blot showing relative levels of TET3, H3K27me3, and total H3 in SK-N-BE2 wild type and *TET2/3* CRISPR-Cas9 mediated knockout cells.

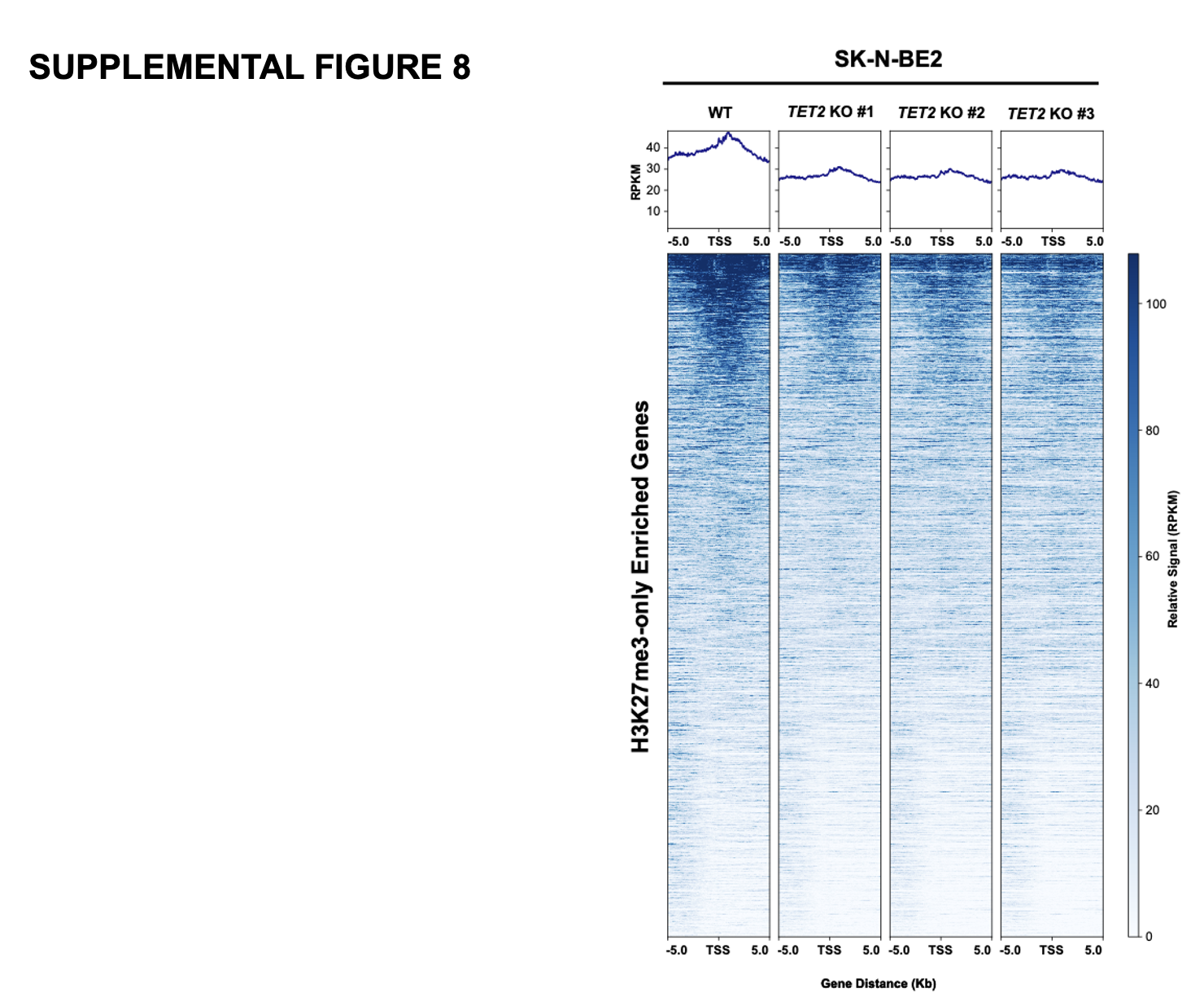

**SUPPLEMENTAL FIGURE 8: Deposition of H3K27me3 on H3K27me3-only enriched genes in WT and *TET2* KO SK-N-BE2.** H3K27me3 profile of the TSS of H3K27me3-only marked genes in WT and *TET2* KO SK-N-BE2 cell lines.

|  | ***TET1* sgRNA (Catalog Number; Viral Titer)** | **DNA target sequence** |
| --- | --- | --- |
| 1 | VSGH11936-248465790; titer: 2.58 x 10^7^ TU/mL | ATCAATATCCCTACCCACAG |
| 2 | VSGH11936-247518826; titer: 4.51 × 10⁷ TU/mL | GCCTTCCAGATTAGTCAGGA |
| 3 | VSGH11936-247566442; titer 2×10⁷ TU/mL | CTGCCATCACGTTAGCACAC |

**SUPPLEMENTAL TABLE 1: *TET1* sgRNAs utilized in the current study.**

**SUPPLEMENTAL TABLE 2: *TET2* sgRNAs utilized in the current study.**

|  | ***TET2* sgRNA (Catalog Number; Viral Titer)** | **DNA target sequence** |
| --- | --- | --- |
| 1 | VSGH111936-247665810; titer: 1.63 x 10^7^ TU/mL | GCAAAGCTCAGTGTTCACTA |
| 2 | VSGH111936-247670218; titer: 2.78 x 10^7^ TU/mL | TAGCATTGCAGCTAGTTTAC |
| 3 | VSGH111936-247797122; titer: 5.33 x 10^7^ TU/mL | GTCATTTGATTGGAGAGATT |

**SUPPLEMENTAL TABLE 3: *TET3* sgRNAs utilized in the current study.**

|  | ***TET3* sgRNA (Catalog Number; Viral Titer)** | **DNA target sequence** |
| --- | --- | --- |
| 1 | VSGH12179-249619287; titer 2.28 × 10⁷TU/mL | TACCAAGCCCAAGGTCAAGG |
| 2 | VSGH12179-250035585; titer: 1.26 × 10⁷ TU/mL | GCAGCTTTCTCCGTTCCCAC |
| 3 | VSGH12179-249635367; titer: 1.89 x 10^7^ TU/mL | GGACAATCTGTCGGACAGGT |

**SUPPLEMENTAL TABLE 4: Combinations of *TET1* and *TET3* sgRNAs utilized to generate *TET1/TET3* double knockout SK-N-BE2 cell lines.**

| **Sample Number** | **Combinations of *TET3* and *TET1* sgRNAs** |
| --- | --- |
| #1 | VSGH12179-249619287 + VSGH11936-248465790 |
| #2 | VSGH12179-250035585 + VSGH11936-248465790 |
| #3 | VSGH12179-249635367 + VSGH11936-248465790 |
| #4 | VSGH12179-249619287 + VSGH11936-247518826 |
| #5 | VSGH12179-250035585 + VSGH11936-247518826 |
| #6 | VSGH12179-249635367 + VSGH11936-247518826 |
| #7 | VSGH12179-249619287 + VSGH11936-247566442 |
| #8 | VSGH12179-250035585 + VSGH11936-247566442 |
| #9 | VSGH12179-249635367 + VSGH11936-247566442 |

**SUPPLEMENTAL TABLE 5: Combinations of *TET2* and *TET3* sgRNAs utilized to**

**generate *TET2/TET3* double knockout SK-N-BE2 cell lines.**

| **Sample Number** | **Combinations of *TET2* and *TET3* sgRNAs** |
| --- | --- |
| #1 | VSGH111936-247670218 + VSGH12179-249635367 |
| #2 | VSGH111936-247665810 + VSGH12179-250035585 |
| #3 | VSGH111936-247670218 + VSGH12179-250035585 |
| #4 | VSGH111936-247665810 + VSGH12179-249635367 |
| #5 | VSGH111936-247797122 + VSGH12179-249635367 |
| #6 | VSGH111936-247797122 + VSGH12179-250035585 |

**SUPPLEMENTAL TABLE 6: PCR primers utilized in the bisulfite sequencing experiments in the current study.**

| **Gene_direction** | **Sequence** |
| --- | --- |
| SLC6A17_F | GTTTGTGATTTTGATAGTAATATTGGTA |
| SLC6A17_R | AAACATAAATACAACAAAAACCTCTC |
| KCNQ3_F | ATATTTTTAGAAGATGTTATAAATGTGT |
| KCNQ3_R | AATATTCCACACTAAAAACTTAAAAAAC |
